# SPRI: Structure-Based Pathogenicity Relationship Identifier for Predicting Effects of Single Missense Variants and Discovery of Higher-Order Cancer Susceptibility Clusters of Mutations

**DOI:** 10.1101/2022.09.27.508720

**Authors:** Boshen Wang, Xue Lei, Wei Tian, Alan Perez-Rathke, Yan-Yuan Tseng, Jie Liang

## Abstract

We report the Structure-based Pathogenicity Relationship Identifier (SPRI), a novel computational tool for accurate evaluation of pathological effects of missense single mutations and prediction of higher-order spatially organized units of mutational clusters. SPRI can effectively extract properties determining pathogenicity encoded in protein structures, and can identify deleterious missense mutations of germ line origin associated with Mendelian diseases, as well as mutations of somatic origin associated with cancer drivers. It compares favorably to other methods in predicting deleterious mutations. Furthermore, SPRI can discover spatially organized pathogenic higher-order spatial clusters (patHOS) of deleterious mutations, including those of low recurrence, and can be used for discovery of candidate cancer driver genes and driver mutations. We further demonstrate that SPRI can take advantage of AlphaFold2 predicted structures and can be deployed for saturation mutation analysis of the whole human proteome.

## 1 Introduction

Whole-genome sequencing and whole-exome sequencing provide powerful means to assess genetic diversity among individuals [1, 2]. Among hundreds of variants in the coding regions [3, 4, 5], missense mutations have the highest frequency of occurrence and can potentially affect molecular functions upon residue substitutions [5, 6, 7, 8]. However, majority of the variants are incidental and have limited effects on protein functions. In addition, missense mutations usually have low recurrence [9, 10, 11, 12], and it is difficult to identify pathological mutations over a noisy background. It also challenging to recognize the collective effects of multiple mutations of low recurrences. It is therefore important to determine whether a missense mutation leads to deleterious phenotypic changes, and whether multiple low recurrence mutations act cooperatively to affect protein function as a higher-order unit.

As functions follow forms, protein structures provide rich information on how missense mutations may affect proteins functions. With rapid progress in structural biology [13, 14, 15, 16] and breakthroughs of AlphaFold2 and RoseTTAFold in structure prediction [17, 18], over 200 millions proteins now have their structures available [17, 19], allowing analysis of the whole protein universe.

However, most current methods to evaluate potential pathological effects of missense mutations are based on sequence alignment and conservation analysis [20, 21, 22, 23, 24, 25, 26], and do not take advantages of structural information [20, 21, 22, 23, 24, 25, 26]. Some require annotated knowledge of functional domains, which is not uniformly available [20] (see Table 1), others exhibit uneven performance on different datasets, indicating lack of transferability [21, 27]. In addition, it is difficult to identify individual mutational events that will lead to diseases: over 70% mutations collected from cancer samples are predicted as deleterious by Polyphen-2 [24], while only a few mutations are believed to drive tumorigenesis and most cancer mutations are passenger mutations [9, 10, 11, 12].

**Table 1:**
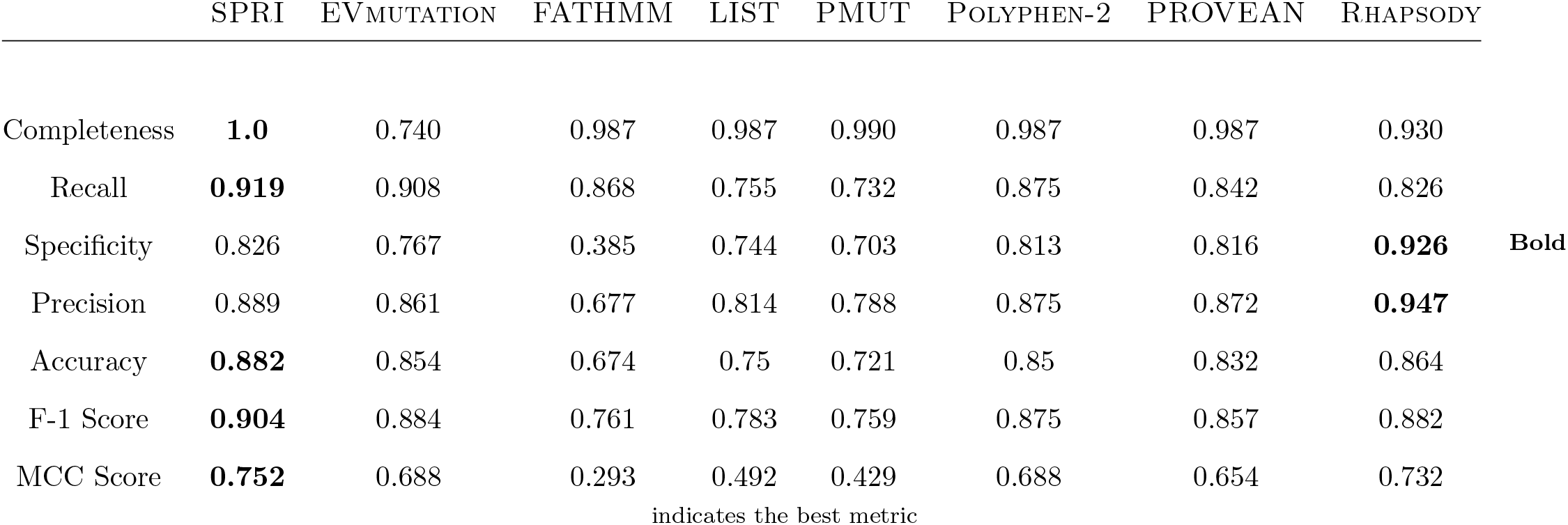
Performance of Predictions on the UnifyPDBFull 5-Fold Test Datasets

Recent methods incorporating protein structural information have improved predictions of deleterious missense mutations [28, 29, 30, 31]. Methods have also been developed that provide mapping of the spatial positions of mutation sites, facilitating identification of clusters of mutation sites close to known functional regions [32, 33, 34, 35, 36, 37]. However, how to extract information encoded in protein structure to improve assessment of pathological effects of missense mutations. It is challenging to determine the relevant factors affecting mutation effects from the myriad of information contained in proteins structures. It is also difficult to discover *de novo* a higher-order spatial units of mutation clusters of low recurrence in a principled fashion without any *a prior* information.

Here we present a new method called SPRI (**S**tructure-Based **P**athogenicity **R**elationship **I**dentifier), which provides quantitative assessment of pathological effects of missense mutations, identifies deleterious mutations, and uncover higher-order structural units of deleterious mutation clusters. Our approach is based on the fact that biochemical functions require specific spatial arrangement of multiple residues and atoms to maintain the structural scaffold and to present a binding or functional surface with the required physico-chemical textures [38]. Mutations altering this microenvironment may affect the execution of biochemical functions. With computation of protein surface pockets [36, 39, 41, 42, 43], novel profiles of interaction at different range, and solvent accessible surface area using alpha shapes [40, 39, 45], we integrate the structural information with evolutionary signals of the mutation site derived from multiple sequence alignment. With further incorporation of changes in biophysical properties upon mutation, a random forest predictor is trained to provide quantitative estimation of pathological effects [46]. The geometric analysis of protein structure and pathognecity estimation further allow us to discover rigorously defined higher-order structural units with enriched accumulation of pathological mutations.

SPRI is among the best in predicting pathogenic missense mutations of Mendelean diseases. Using benchmark data with reduced inherent bias of uneven representations of neutral and deleterious mutations, our methods outperforms several state-of-the-art methods using a number of quality metrics (AU-ROC, AU-PRC, MCC, and F-1 score) [20, 21, 22, 23, 24, 25, 29].

SPRI can also predict cancer driver mutations of somatic origin with high sensitivity. As our method can captures essential properties required for proper biochemical functions, its effectiveness in identifying disease-causing mutations is transferable. We show SPRI can effectively evaluate pathogenicity of mutations of somatic origin and can identify cancer driver mutations. These predictions are made without re-training using different datasets or additional parameter adjustment. This is in contrast to current methods, which make inconsistent predictions with higher rate of false negative predictions [21, 29].

SPRI can discover spatially organized pathogenic higher-order spatial (patHOS) units of deleterious mutations, including those of low recurrence. In a large scale analysis of 239,164 cancer mutations on 2,712 human proteins, we discovered 567 patHOS units in 189 genes, each harboring a significant number (*≥*5) of deleterious mutations, indicative of their high susceptibility to cancer. Among these, 83.7 % of the mutations have *<* 3 recurrence count per each mutation. Furthermore, we uncover the 161 non-CGC genes containing patHOS clusters, 29 of which are strong candidates of cancer driver genes.

In addition, SPRI can be used for *in silico* saturation mutagenesis and can predict pathological effects of systematic mutations at each site of a protein. Finally, SPRI can be deployed for global studies of mutation effects of the whole protein universe. Using a set of proteins with AlphaFold2-predicted structures, we show SPRI can make reliable predictions on the pathological effects of missense mutations at sites whose structure are predicted at high or moderate confidence level.

Overall, we expect that SPRI can be applied broadly to assess pathological effects for mutations regardless of the disease types. It can also be used to discover higher-order mutation clusters, and to provide *in silico* saturation mutagenesis analysis. It can be applied proteome-wide scale studies using known and predicted protein structures.

## 2 MATERIAL AND METHODS

### 2.1 Benchmark Dataset with Reduced Bias

For reliable prediction of pathological effects and deleterious mutations, a dataset that accurately reflects the natural landscape of mutations is critical. Existing benchmark datasets such as HumVar, HumDiv, ExoVar, VariBench, and PredictSNP [24, 22, 47, 48, 50] are biased, as deleterious mutations and neutral mutations are often from different proteins, which may have different degree of deleteriousness depending on the number of functional domains, size of the functional surfaces, and other factors (see details in Supplemental Information). Furthermore, many proteins were not fully investigated, thus variants lacking annotation of pathological effects may be simply a reflection of lack of knowledge, rather than positive information about these mutations being neutral. As only a small proportion of proteins possess both deleterious and neutral variants (see SI) in these datasets, inconsistency in predictions when applied to different data may result, suggesting that improved training data are needed for robustness and transferability.

We construct two data sets based on the HumDiv, HumVar and PredictSNP datasets as benchmarks [24, 50]. To ensure the same level of knowledge of both types of mutations, we select only proteins with annotations of both deleterious and neutral mutations, and have high-quality structures. The UnifyPDBFull dataset contains 4,231 deleterious variants and 2,791 neutral variants, which are derived from 252 single-chain proteins with full structural coverage. The UnifyPDBAcceptable dataset contains 5,999 deleterious variants and 3,485 neutral variants, which are derived from 377 proteins with 444 polypeptide chains with full and partial structural coverage, including the UnifyPDBFull set as a proper subset. Our datasets are smaller but have much less bias, providing more accurate representations of the landscape of deleterious and neutral mutations in proteins.

### 2.2 Structural, Biophysical, and Evolutionary Properties

#### 2.2.1 Protein Surface Pocket and Geometric Location of Mutated Residues

Surface pockets on proteins provide the local microenvironment for ligand binding and biochemical reactions [38, 51], while core residues may contribute to folding stability [44]. We therefore classify each residue into the categories of surface pockets residues, interior buried residues, and residues on other surface region [41, 45]. We also calculate the solvent accessible surface area of each residue. Pocket construction, surface and buried core residues, as well as solvent accessible surface areas are computed using the alpha shape method and are accessible from the Computer Atlas of Surface Topography of Proteins (CASTp) server [36, 52].

#### 2.2.2 Contact Profiles of Short Range Atomic Interactions

An amino acid residue plays its biochemical roles in conjunction with its spatial neighbors as part of a favorable microenvironment for reactions. While residue distances have been used to describe the local environment [31, 53], complex patterns in atomic interactions and side-chain packing arrangement are difficult to capture by distance-based profiles.

To capture immediate atomic interactions at the mutation site, we construct its atomic contact profiles. We obtain the contact network from the weighted alpha shape, which corresponds to the Voronoi diagram of the protein structure with non-empty nearest neighbor atomic contacts. As only those with physical contacts are found in alpha shapes [54, 55], this allows us to focus on exact atomic contacts without the distraction of noise inherent in contacts defined by distances.

Specifically, we first computed the weighted Delanuay triangulation of the atoms of the protein structure. We then obtain its alpha shape by generating a filtration of the Delaunay simplicial complex [56, 57] We further select alpha edges connecting different residues and obtain inter-residue atomic contact network, where nodes are atoms and alpha edges connect nearest-neighboring atoms whose volume overlaps.

We consider only four atom types of C, N, O, and S, distinguishing atoms provided by the mutated residue and its neighboring residue. Altogether, we have 16 types of element pairs for atomic interaction.

#### 2.2.3 Interaction Profiles of Residues of Intermediate and Long Range

Intermediate and long range interactions can influence the microenvironment of the mutation site residue via electrostatic interactions and indirect steric effects [58, 59, 60]. We construct interaction profiles of intermediate- and long-range at the residue level to account for such effects. Interacting residues are again identified by the atomic connection network derived from alpha shape.

Specifically, we start with residues in direct atom contacts with the mutation site, as recorded in the short-range contact profile. From these *first layer* contact residues. We use a Breadth-First Search (BFS) algorithm to identify the next layer of residues in atomic interactions with them [61]. These are residues of *intermediate-range* interactions. We apply BFS again to identify the next layer of partner residues through atomic contacts. These newly identified residues are in *long-range* interactions with the mutation site. We construct the intermediate- and long-range interaction profiles, each a 20-dimensional integer vector recording the number and type of residues interacting with the mutation site.

#### 2.2.4 Higher-Order Spatial (HOS) Cluster

Here we define <u>h</u>igher-<u>o</u>rder spatial (HOS) cluster rigorously based on the computed alpha shape. We take the mutation site residue as the anchor, and incorporate all residues with short-, intermediate-, and long-range interactions from the anchor as its HOS cluster. HOS clusters provide an *a priori* defined spatial units from structures for analysis of higher-order effects of mutations.

#### 2.2.5 Salt Bridge Interactions

Salt bridge formed between ion pairs can stabilize protein conformation and provide protons for catalytic reactions [64]. We count the total number of salt bridges that an ionizable residue participates in. The criterion is that the sidechain oxygen atoms in an acidic residue and the side-chain nitrogen atoms in a basic residue has a distance *≤* 3.2 Å.

#### 2.2.6 Changes in Biophysical Properties in Variants

Different substitutions occurring at the same site may have dramatically different effects. It is therefore important to consider changes in biophysical properties upon a substitution to obtain finer distinction of mutation effects of different substitution patterns at a given mutation site. We record differences in each atomic chemical type upon mutation. If Gly is involved, we record the change as *C_β_*. We further calculate changes in charges upon mutations.

#### 2.2.7 Robustness of Structural Properties

The structured-derived properties are robust. Using 5 different PDB structures of the same protein uroporphyrinogen decarboxylase, about 85% of structure-derived features are of the same value or small deviations, leading to the same prediction results. (see SI for more details.)

#### 2.2.8 Evolutionary Signals

We use Blosum62 matrix to capture general patterns of substitutions [63]. To incorporate protein-specific evolutionary information, we construct multiple sequence alignments (MSA) using the full protein sequence instead of domains [62]. We take homologs with full-length sequence identity *≥* 30% in the reference proteomes of other eukaryotic species, from which we construct the MSA using CLUSTAL OMEGA [62]. Overall, 94.4% protein-specific MSAs analyzed in this study (356 out of the 377 UnifyPDBAcceptable proteins) have adequate alignment depth of *≥* 100, ensuring that reliable evolutionary signals can be detected. We then calculate the entropy value of the mutation site using the aligned MSA [62]. In addition, frequencies of occurrence of the wild-type and the mutated-type residues in the MSA for each mutation site are also recorded. More details can be found in SI.

### 2.3 Predicting Pathological Effects

#### 2.3.1 Pathogenicity of Missense Mutation

We integrate all information extracted from structure and sequence and generate a probability score measuring the pathological effects of a substitution. We construct a random forest classifier [65] as it is robust from overfitting [46]. The prediction score *π*_del_ is calculated from the ratio of the number of trees supporting the hypothesis that the variant is pathological against the total number of trees in the forest [46]. The overall architecture of our method is shown in Figure 1.

**Figure 1:**
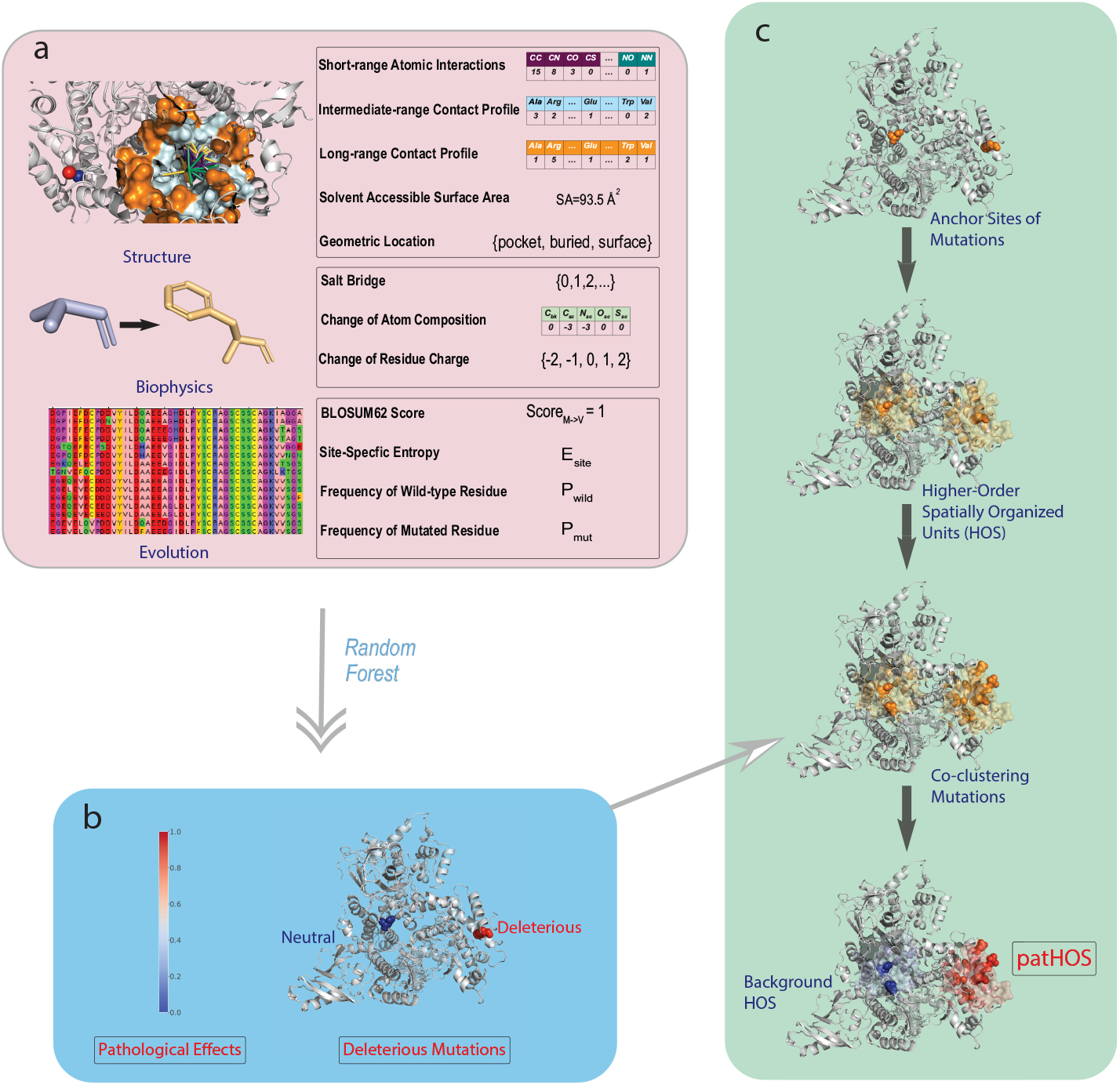
Overview of the SPRI method. a) SPRI utilizes three types of features based on protein structures, biophysical properties, and evolutionary signals, respectively. Structural features include the atomic and residue interaction profiles computed using alpha shapes, which capture the site-specific micro-environment of the mutation site. Geometric shape at the mutation site and at neighboring residues, as well as solvent accessible surface area, are also incorporated. Features of biophysical properties include possible salt bridges formed by ionizable residues, changes in charge upon substitutions, as well as changes in side-chain and backbone atoms. To obtain features of evolutionary signals, a knowledge database containing *∼*24 millions protein sequences from reference proteomes of eukaryotic species in UniProt is built where homologs are identified, so multiple sequence alignment can be constructed to obtain sitespecific metrics of entropy, frequencies of wild-type and mutated residues. b) These features, except geometric location, are vectorized into numerical values. A random forest classifier is then trained to generate probability measure of the likelihood of pathological effect, from which a binary classification is made. c) SPRI can discover pathogenic higher-order spatial clusters (patHOS) of deleterious mutations. For each observed mutation site, SPRI constructs its higherorder structural unit (HOS) based on geometric computation, and evaluate the overall pathogenecity of all mutations located in the HOS. After normalization by the size of HOS, those clusters with (*≥*5) deleterious mutations and pathogenecity score *≥ θ* = 0.05 are classified as pathogenic HOS (patHOS) and are distinguished from clusters with background mutation.

#### 2.3.2 Pathogenicity of Higher-Order Spatial Cluster

The degree of pathogenicity of a HOS cluster is calculated by summing up the pathology probability scores of its residues, which is then normalized by the cluster size. We take each the site of each mutation as the anchor and calculate the associated cluster pathogenicity score. Those above a threshold of *θ* = 0.05 and harbor *≥*5 deleterious mutations are considered to be pathogenic high-order spatial cluster (patHOS).

Additional details can be found in the SI.

## 3 RESULTS

### 3.1 SPRI Can Capture Functional Spatial Patterns without Prior Information

**Figure 2:**
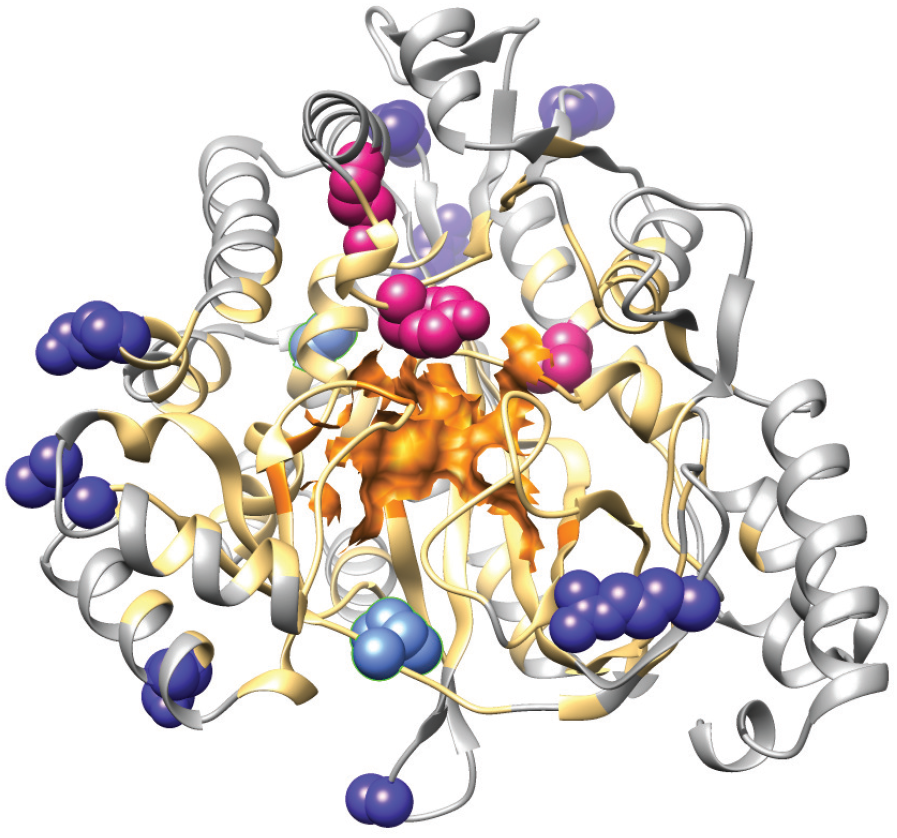
Illustration of different structural regions and mutation sites of the human glutathione synthetase, which are color-coded: the catalytic region (orange), its neighboring environment (light orange), outside region (grey), the correctly predicted deleterious mutations (pink), the correctly predicted neutral mutations (dark blue), and the false predictions of deleterious mutations (light blue). All 4 deleterious mutation site residues (L188P, Y270C, Y270H, R283C) are located within the neighboring environment of the catalytic region, and are correctly predicted. Eight correctly predicted neutral predictions are located in the outside region, and the two false deleterious predictions are located at the boundary of the catalytic neighboring environment.

We use human glutathione synthetase (GSS, UniProt: P48637, PDB: 2HGS chain:A) for illustration. This protein has 524 residues. There are 4 known deleterious variants occurring at 3 mutation sites. Among these, the variant R283C leads to glutathione synthetase deficiency symptom [66]. There are 10 neutral variants occurring at 10 different mutation sites. Our method provides reliable predictions on these variants without specification of the catalytic residues or any other prior annotations: All 4 deleterious variants (L188P, Y270C, Y270H, R283C) are correctly predicted, 8 out of 10 neutral variants (K95E, A134T, P202T, H290C, V343M, R418Q, Q435H, E353K) are correctly predicted, and only 2 neutral variants (S80N, I401T) are incorrectly predicted to be deleterious.

Post-prediction analysis of the results show that SPRI can extract complex spatial patterns contributing to functions without *a priori* knowledge. The catalytic region of GSS has 9 residues, with E144, N146 and E368 binding to magnesium, R125, S151 stabilizing the tetrahedral intermediate, K305, K364, G369 and R450 stabilizing the phosphate substrate and the leaving groups electrostatically [68]. We take these 9 residues and compute the 3-layer interaction profiles using alpha shapes. Residues identified form the *neighbor environment* of the catalytic region. The rest of the residues are regarded as *outside residues*. We found that all deleterious variants are located within the catalytic region and its neighbor environment, and the 8 neutral variants with correct predictions are all outside residues. The two incorrectly predicted neutral variants are at the boundary of the neighbor environment.

SPRI’s ability in extracting spatial patterns important for functions from structures are general. We have further examined a total of 19 enzymes whose biochemistry are well understood and catalytic residues well annotated in the M-CSA enzyme database [69]. Statistical test over these enzymes show that deleterious variants are more preferably located at the catalytic or its neighboring region, while neutral variants are more likely to occur at the outside region (*p* = 4.2 *×* 10*^−^*^14^ by Fisher’s exact test. It is important to note that SPRI does not require any prior knowledge on enzyme catalytic residues for predictions.

### 3.2 SPRI Is Highly Effective in Predicting Pathogenic Effects of Mendelian Mutations

To evaluate the effectiveness of SPRI in predicting deleterious Mendelian mutation, we use stratified *k*-fold cross-validations on both the UnifyPDB-Full and the UnifyPDBAcceptable datasets [70]. We focus on the former as it contains proteins whose structures cover the full sequences, ensuring accurate representations of structural properties. We compare our results with those from the methods of EVmutation, FATHMM, LIST, PMut, Polyphen-2, PROVEAN and Rhapsody [20, 21, 22, 23, 24, 25, 29]. We report completeness, recall, specificity, precision, accuracy, F-1 score, and MCC for comprehensiveness. Among these metrics, MCC is an informative and balanced metric to symmetrically evaluates both groups of positives and negatives with adjustment of ratio of subgroups [71].

The 5-fold average performance of our method on UnifyPDBFull stratified test data has an F-1 score and MCC of 0.904 and of 0.752, respectively, which are the highest among all 8 methods compared (Table 1). Our method also has the highest accuracy (0.882) and highest recall (0.919), indicating it is highly sensitive in identifying deleterious mutations. In addition, our method has an excellent specificity of 0.826 in identifying neutral mutations correctly, which are better than all other methods except Rhapsody (0.926). Our methods outperforms Rhapsody in several important regards: Rhapsody has a lower recall of 0.826, and it can make predictions on less variants in UnifyPDBFull (0.930), whereas our SPRI method has a higher recall of 0.919 and makes predictions on all variants.

The Receiver Operating Characteristic (ROC) and the Precision-Recall (PR) curves of the overall performance of prediction at different thresholds are shown in Figure 3. On the stratified UnifyPDBFull test dataset, our method has an AU-ROC of 0.946 and an AU-PRC of 0.966, which are the best among all methods. These results show that our method is robust and performs well at different thresholds. Results using cross-validation data of (UnifyPDBAcceptable) are similar to that of UnifyPDBFull, where our method has the best performance when measured by the metrics of F1-score, MCC value, accuracy, and AUCs of ROC and PR curves (Table 2 and SI).

**Figure 3:**
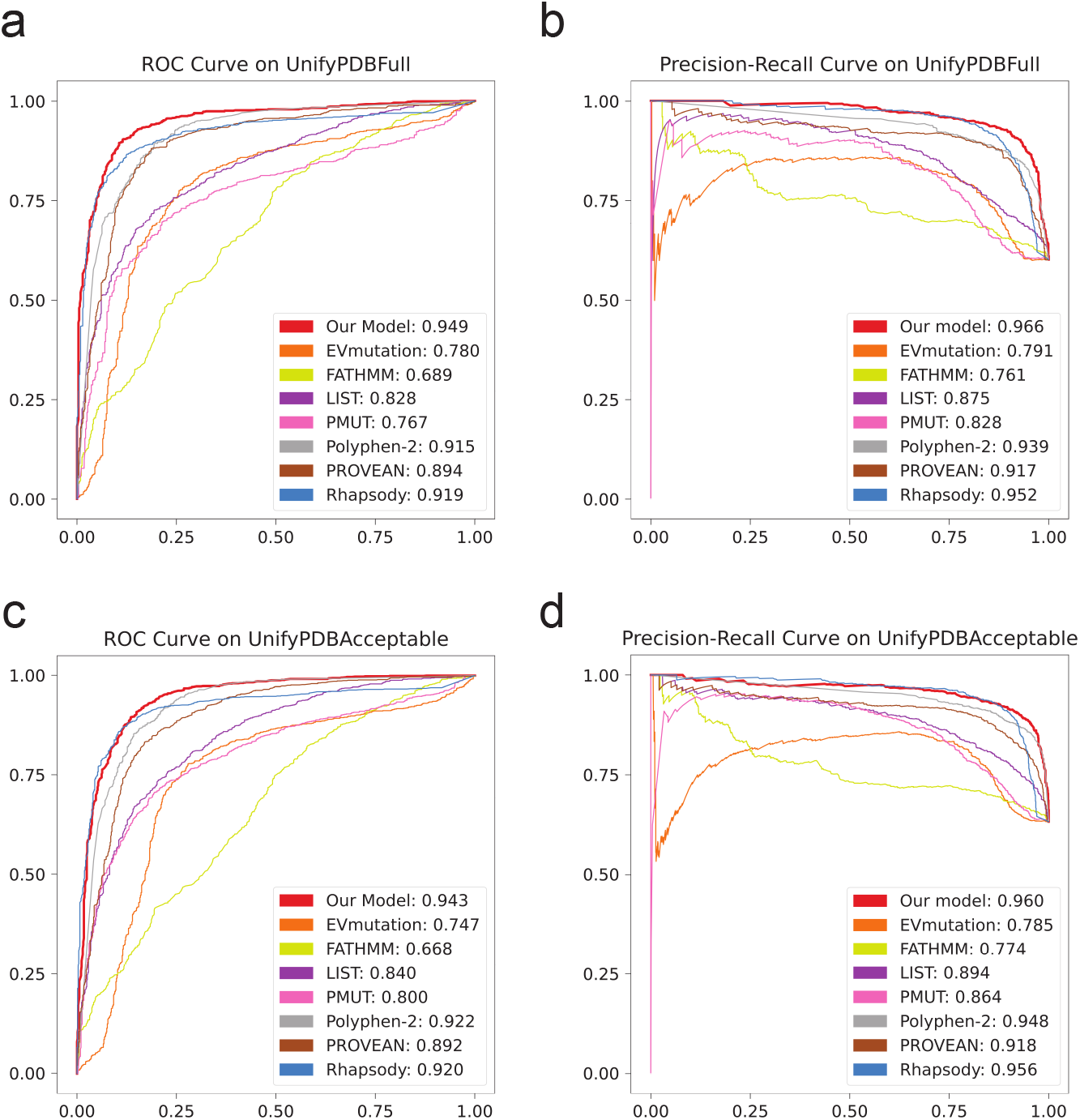
Receiver Operating Characteristic (ROC) curves and Precision-Recall (PR) curves of testing results using SPRI and other methods on the UnifyPDBFull and UnifyPDBAcceptable test datasets. (a) The ROC curves and their AU-ROCs using the UnifyPDBFull test dataset. SPRI has the highest AU-ROC of 0.949. (b) The PR curves and their AU-PRCs using the UnifyPDBFull test dataset. SPRI has the highest AU-PRC of 0.966. (c) The ROC curves and their AU-ROCs using the UnifyPDBAcceptable test dataset. SPRI has the highest AU-ROC of 0.943. (d) The PR curves and their AU-PRCs using the UnifyPDBAcceptable test dataset. SPRI has the highest AU-PRC of 0.960.

**Table 2:** Performance on the UnifyPDBAcceptable 5-Fold Test Datasets

|  | SPRI | EV<sub>MUTATION</sub> | FATHMM | LIST | PMUT | POLYPHEN-2 | PROVEAN | RHAPSODY |
| --- | --- | --- | --- | --- | --- | --- | --- | --- |
| Completeness | **1.0** | 0.711 | 0.990 | 0.990 | 0.987 | 0.990 | 0.990 | 0.941 |
| Recall | **0.913** | 0.902 | 0.846 | 0.753 | 0.747 | 0.881 | 0.824 | 0.822 |
| Specificity | 0.828 | 0.766 | 0.405 | 0.766 | 0.732 | 0.809 | 0.817 | **0.925** |
| Precision | 0.902 | 0.878 | 0.707 | 0.846 | 0.829 | 0.888 | 0.884 | **0.952** |
| Accuracy | **0.882** | 0.854 | 0.683 | 0.758 | 0.742 | 0.855 | 0.821 | 0.859 |
| F-1 Score | **0.907** | 0.890 | 0.771 | 0.797 | 0.786 | 0.884 | 0.853 | 0.882 |
| MCC Score | **0.744** | 0.676 | 0.281 | 0.505 | 0.466 | 0.689 | 0.628 | 0.721 |

indicates the best metric

Overall, SPRI has the best performance by most measurements among the 8 methods compared, and is robust regarless of the thresholds, despite the fact that SPRI is derived on a training data set of much smaller size.

### 3.3 Features Collectively Contributes to Model Performance

To understand the relative contributions to effectiveness of predictions from different types of features, we compare the model trained on the full set of features with models trained on different subsets of features. Specifically, we trained additional models with evolutionary only, structural+biophysical only(non-evolutionary), structural only, and biophysical features only.

We compare the average values of AUROCs and AUPRCs of models based on 5-fold cross-validation of the UnifyPDBFull test datasets (SI Table 4 and SI Figure 3-7). Among these, the model trained with structural+biophysical (non-evolutionary) features has the best AUPRC (0.917) and the second-best AUROC (0.879). The model trained on evolutionary features has comparable AUPRC (0.913) and AUROC (0.891).

These results show that structural+biophsycial features contribute equivalently as evolutionary signals to the model performance, even though each individual structural and biophysical feature has lower feature-wise importance according to mean decrease in the Gini coefficient (SI Figure 2). It is notable that none of the subsets of features is capable of achieving comparable performance with the full set of features, suggesting that structural, biophysical, and evolutionary complement each other, and collectively contribute to the overall performance of the full model. Detailed information can be found in the SI 4.

Analysis of Feature Importance.

### 3.4 SPRI Can Predict Cancer Driver Mutations of Somatic Origin

**Figure 4:**
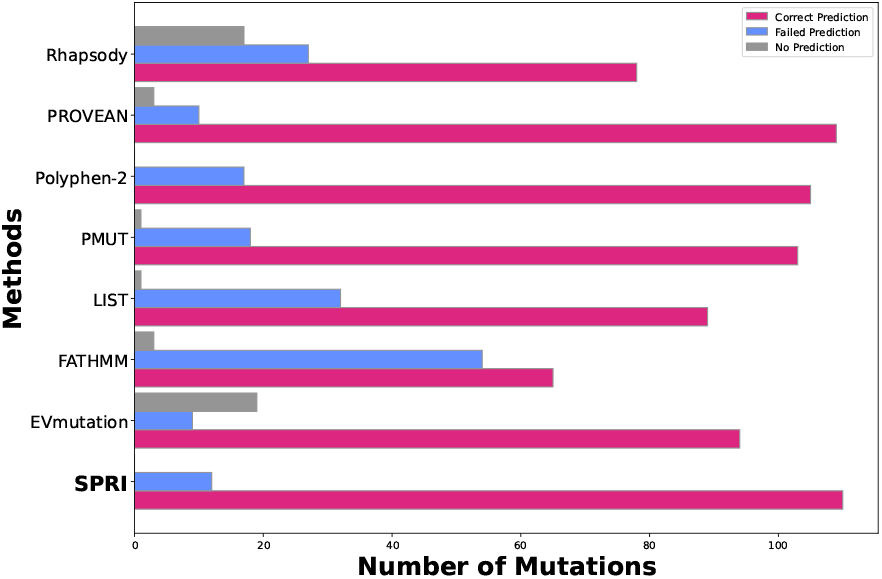
Predictions of cancer driver mutations among the set of cancer census mutations (cmc) tier-1 mutations. For each method, the numbers of correct prediction (pink), failed prediction (light blue), and missing predictions (gray) are listed. Overall, our SPRI method has the highest number of 110 correct predictions out of 122 driver mutations.

Somatic mutations provide one of the most important means to trigger abnormal cell growth that may lead to tumorigenesis [72, 73, 74]. Identifying somatic mutations that drive cancer development can facilitate development of therapeutics targeting these mutated proteins [75]. However, there are millions of missense mutations accumulated in cancer patients, the majority of which exhibit low recurrence [6]. Current methods based on frequency counting can provide gene-level predictions, but are challenged in identifying local structure regions with higher pathogenicity, or in pinpointing individual cancer driver mutations [9].

While SPRI has been developed and evaluated using mutations of Mendelian disorders, we hypothesize that SPRI captures the essential characteristics of pathological mutations and the predictive model is transferable to predict cancer somatic mutations. To test this, we take the recently available annotations of tier 1 cancer census mutation from the COSMIC v92 as the ground truth [6], as there is strong evidence for their roles in tumorigenesis from experimental, clinical, and *in silico* analysis [6]. This dataset contains 201 missense mutations with both wild-type residue and substituted amino acid. Among these, 122 mutations can be mapped to 50 mutated residues on 26 proteins with known structures. We build a separate predictor following the same procedure but with the 26 cancer relevant proteins removed from the training process.

Our method has excellent performance in identifying cancer driver mutations. It can correctly classify 110 mutations out of the 122 structure-mapped mutations as deleterious. We also tested the methods of PROVEAN, Polyphen2 and PMut, which have comparable performance, identifying 109, 105, and 103 deleterious mutations, respectively. EVmutation identifies 94 deleterious mutations, with 9 incorrect predictions and 19 missing predictions. Rhapsody identifies 78 out of the 122 deleterious mutations, at a significantly reduced level of effectiveness than Mendelian disorders. FATHMM identifies 65 deleterious mutations, with 54 incorrect predictions and 3 missing predictions. Overall, our method has excellent performance in identifying confirmed cancer driver mutations.

### 3.5 SPRI Can Distinguish Higher-Order Structural Clusters of Low Recurrence Mutations But Strong Cancer Susceptibility from Background Clusters

By evaluating the pathogenicities of the higher-order spatial (HOS) clusters for each known mutation site taken as the anchor, we can distinguish those with strong disease susceptibility from background clusters. As an illustration, two HOS clusters with different levels of pathogenicities exist on the protein XPD (encoded by ERCC2, Figure 5b). The HOS cluster centering at I668 has 77 residues in total, harboring 8 deleterious mutations (T460A, T480A, T484A, T484R, G607A, G665A, G665R, and I668F), all are of low recurrence (1 for 7 mutations and 3 for 1 mutation). With a cluster pathogenicity score of 0.078 *> θ*=0.05 and *≥*5 deleterious mutations, this cluster is classified as a pathogenic HOS (patHOS) cluster with strong disease susceptibility. Indeed, this patHOS cluster contains the binding sites to ATP and a portion of the functional domain mediating interaction with MMS19. The accumulation of multiple low recurrence mutations with deleterious effects in this cluster likely strong affect the function of this protein.

**Figure 5:**
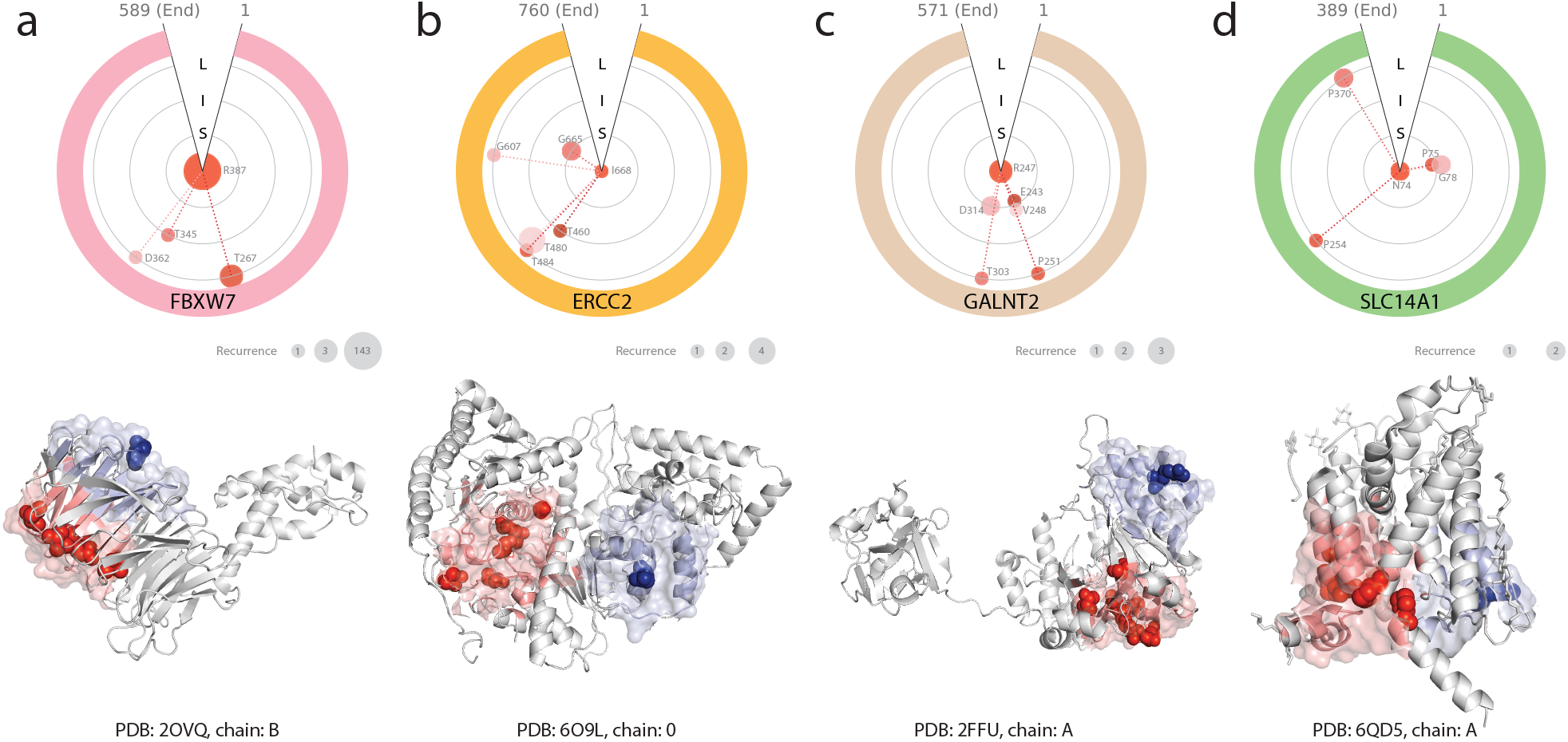
Discovery of pathogenic higher-order structural clusters (patHOS) enriched with low-recurrence deleterious mutations. (top) The polar plots illustrate the spatial relationships between co-clustering deleterious mutation sites and the anchor residue in the same higher-order spatial (HOS) cluster. Mutation sites are clockwise ordered according to their linear sequence index. Light-gray rings with different radii reflect relative distances to the anchor, with **S**, **I**, and **L** for short-, intermediate-, and long-range distance, respectively. The size of the colored disks are proportional to the recurrence counts in the pan-cancer data, except R387 of FBXW7. The recurrence count numbers are listed in the grey disks. (bottom) The corresponding protein structures with the mutational paTHOS clusters highlighted in light red. The mutation sites with deleterious effects are shown as red spheres. For comparison, background spatial structural clusters containing only neutral mutations are shown in light blue, with the mutation sites shown as blue spheres. a. the patHOS cluster anchoring at R387 in the confirmed cancer gene FBXW7 and a background cluster anchored at E353. b. the patHOS cluster anchoring at I668 in the confirmed cancer gene ERCC2 and a background cluster anchored at Q187. c. the patHOS cluster anchoring at R247 in the novel identified cancer gene GALNT2 and a background cluster anchored at R113. d. the patHOS cluster anchoring at N74 in the novel identified cancer gene SLC14A1 and a background cluster anchored at W171.

In contrast, the HOS cluster centering at Q187 has 37 residues in total, but harboring only 2 neutral mutations (Q187R and Q187K), with a pathogenicity score of 0.006, and is therefore classified as a background cluster. This cluster of residues is likely of low cancer susceptibility.

### 3.6 SPRI Can Discover Novel Candidate Cancer Driver Genes

We applied SPRI to a large scale study to search for candidate cancer driver genes. Using the pan-cancer mutation data of whole genome and whole exome sequencing from COSMIC database (v95), we analyze 239,164 mutations from 200,402 unique mutation site residues on 2,712 proteins with full-coverage PDB structures. We exhaustively take each unique mutation site as the anchor, and calculate the pathogenicity of its HOS cluster.

Over 96.4% of the HOS clusters are found to harbor no more than 3 mutations. Using the threshold of pathogenicity score *≥ θ* = 0.05 and *≥*5 deleterious mutations, we are able to identify 567 patHOS clusters, which likely have strong cancer susceptibility. These clusters are concentrated on cancer driver genes: 200 high-susceptibility patHOS clusters are found on 28 of the 105 known cancer driver genes according to Cancer Gene Census (CGC) [6]. Another 367 clusters are found on 161 of the 2,607 genes not known to be cancer genes (*i.e.*, those are part of the non-CGC genes). Among these 161 genes, we predict that 29 genes to have high likelihood to be cancer driver genes, as each has at least 2 non-overlapping patHOS mutational clusters.

### 3.7 SPRI for In Silico Saturation Mutation Analysis

As SPRI can predict deleterious mutations contributing to both Mendelian disorders and complex diseases such as cancer, it can be used to assess general effects of substitutions of arbitrary sites from the wild type to any of the other 19 amino acid types. Fig 6 depicts an example, where we show the heatmap of pathological effects of all different substitutions at all positions along the full protein sequence of the copper transport protein. The color intensity of each square in the heatmap represents the level of pathological effect of a substitution. As SPRI is scalable, such heatmaps can be computed exome/proteome wide for all proteins with structures.

**Figure 6:**
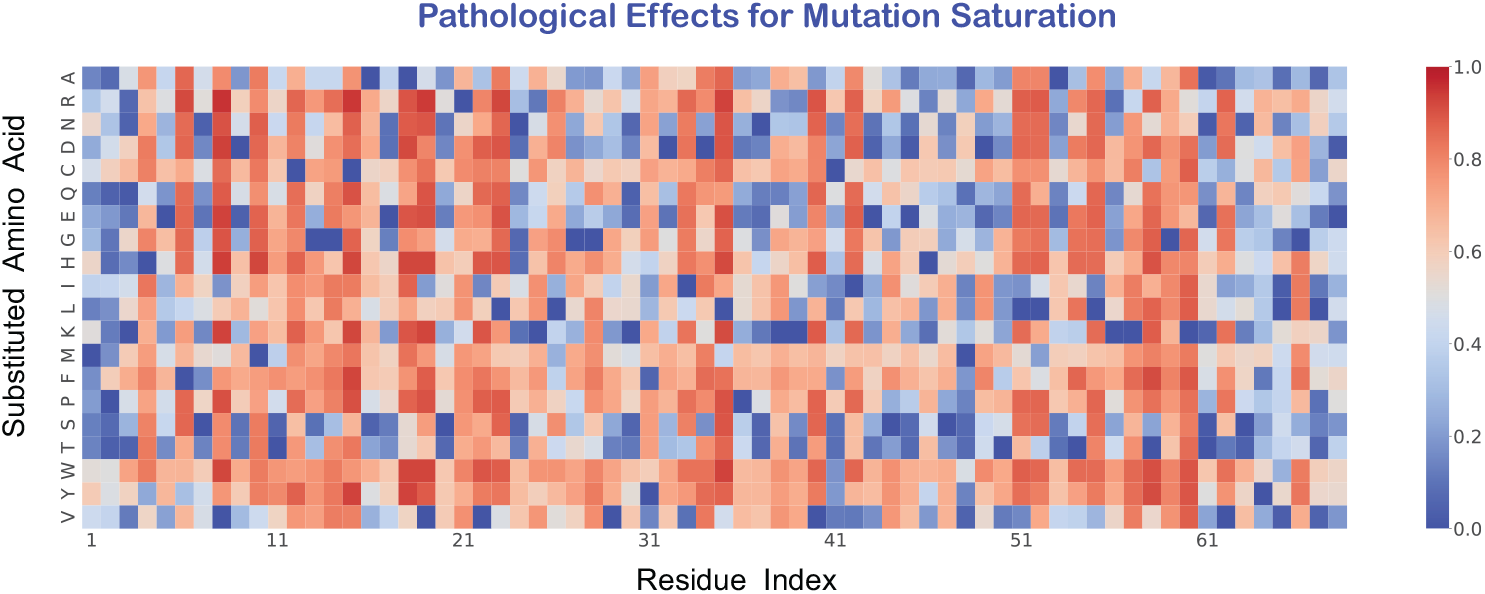
The heatmap of predicted pathological effects of mutation saturation of copper transport protein ATOX1 (Uniprot: O00244). The horizontal axis are the residue index along the sequence, and the vertical axis represents the substituted amino acid types. Each square is color coded by the effects of mutation, where the color intensity encodes the likelihood of pathological effect. A red substitution represents a mutation predicted to be likely a pathological mutation, whereas a blue substitution likely a neutral mutation.

### 3.8 SPRI Can Make Reliable Predictions Using **AlphaFold2 Structures**

SPRI is a structure-based method. As about 200 million proteins structures can be predicted by AlphaFold2 [17, 19], we apply SPRI to a set of proteins with AlphaFold2 structures but no experimental structures and evaluate how well pathological effects of missense mutations can be predicted.

We adopt the same dataset construction procedure as UnifyPDBFull, and take the subset of proteins from HumDiv, HumVar and PredictSNP that have AlaphFold2 structures but no PDB structures, and generate a new dataset UnifyAF2. It contains 4,980 deleterious and 2,600 neutral variants from 422 proteins. We use exactly the same model trained on UnifyPDBFull to predict pathological effects of missense mutations in UnifyAF2, which has no overlap with UnifyPDBFull. We stratify mutation site residues into three categories of high, medium, and low-confidence level of their predicted structures using the pLDDT score from AlaphFold2 (see SI).

We report the ROC curves and PR curves of prediction results in Figure 7. For the high-, moderate-, and low-confidence mutation site residues, our method achieves AU-ROCs of 0.908, 0.905, and 0.854, respectively. The corresponding AU-PRCs are 0.956, 0.950, and 0.810, respectively. Overall, except residues of low-confidence, SPRI can provide reliable assessment of mutation effects using AlphaFold2 structures.

**Figure 7:**
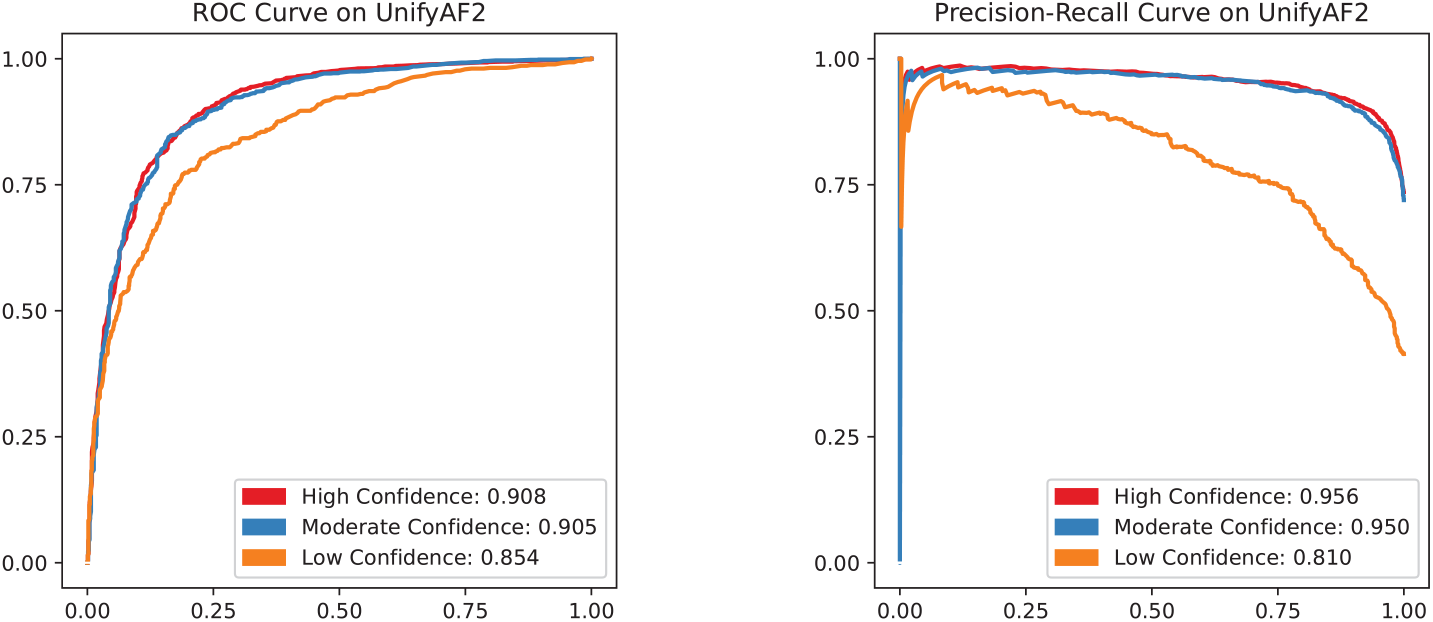
Receiver Operating Characteristic (ROC) curves and Precision-Recall (PR) curves of prediction results using SPRI on the UnifyAF2 dataset of AlphaFold2 predicted structures. The performance is evaluated separately by the AlphaFold2 confidence level of the mutation site residues. (a) The ROC curves and their AU-ROCs using the UnifyAF2 dataset. (b) The PR curves and their AU-PRCs using the UnifyAF2 dataset. Red, blue, and orange curves represent residues with high-, moderate-, and low-confidence pLDDT prediction quality, respectively.

These results indicate that our method can be applied broadly to Alaph- Fold2 predicted structures with reliable performance. As the high- and moderate-confidence residues constitute 61.7% of the 14,850,403 residues among 23,391 AlaphFold2 predicted human protein structures, our method can be applied at whole-exome or whole-proteome scale.

## 4 DISCUSSION

Biochemical functions require specific spatial arrangement of multiple residues and atoms to collectively provide the structural scaffold and reaction surfaces [38, 41, 39]. SPRI can recognize such spatial arrangements through explicit computations of surface pockets, atom and residue-level interaction profiles, and further incorporation of changes in biophysical properties upon mutations. As shown in the case of glutathione synthetase, deleterious variants are likely to occur in the catalytic region and its neighborhood, while neutral variants likely to be outside these regions. With exact geometric computation, these regions can be precisely defined using surface analysis and atomic-residue interactions profiles.

Furthermore, SPRI take advantage of information contained in full structures, and infers pathogenicity automatically without requiring *a priori* knowledge of annotated functional domains or residues. This is different from existing methods where analysis is carried out at the domain level, requiring prior knowledge on clearly defined functional domains as well as demanding significant alignment depth [20]. As a result, SPRI can be applied to a much large number of mutations sites that have no domain annotations. In addition, there is no need of *ad-hoc* re-weighting or other post-processings, unlike the practice of several other methods [22, 24].

SPRI can identify pathogenic higher-order spatially organized units (patHOS) of local regions on protein structures. These units are enriched with strongly deleterious mutations that are well-separated in sequence (as shown in Fig 5). As the structural scaffold and reaction surfaces of biological functions consists of many residues [67, 49], there are higher-order spatially organized clusters of functionally important residues. Our results show that when these are enriched with deleterious mutations, they can exhibit strong pathogenicity, even though individual mutations may be of low recurrence and these PATHOS clusters can elude detection by frequency analysis of sequences. With this approach, we reported the discovery of candidate cancer driver genes and associated PATHOS clusters of mutations.

SPRI has an important advantage over other methods in identifying structural clusters of mutations [33, 37]. SPRI makes the important distinction between a pathogenic higher-order spatially organized unit (patHOS) harboring strongly enriched deleterious mutations and a background structural higherorder spatial (HOS) cluster. No such distinctions are made in methods such as Mutation3D and AlphaCluster [33, 37]. SPRI can then predict cancer driver genes based on the detection of patHOS. While both SPRI and mutPanning, a frequency-based method, can identify all 28 CGC genes as cancer driver genes, no predictions on 160 out of 161 non-CGC genes with strong patHOS identified by SPRI are predicted to have cancer susceptibility by mutPanning [9]. Another important advantage of SPRI is it can pinpoint individual residues where predicted cancer pathogenecity concentrate. No information at residue level was provided by mutPanning, as its prediction is based on simulated mutation rate of the whole gene. Overall, our results show that SPRI can discover higher-order pathological clusters of low-recurrence mutations and distinguish them from otherwise noisy background. Such a task is not possible with prediction methods that works at the gene-level [9, 77].

Our results also sheds some light on an important question, namely, whether pathological missense mutations of different varieties are organized by the same principles, regardless of whether it is of germline or somatic origin, and whether it manifests as Mendelian diseases or complex diseases. Some studies have built pathogenicity models that are disease-specific or dataset-specific. For example, FATHMM has different predictors for inherited disease, cancer, and other specific pathologies [21]. PMut allows user-input to customize training datasets so different predictors for different diseases can be constructed [23]. There are drawbacks with these approaches. FATHMM has very different answers in evaluating pathogenic mutations, depending on whether a predictor for inherited diseases or a predictor for cancer is used. This is likely due to the different level of available annotated information and the necessary adjustment of the model parameter [21]. While Rhapsody works well in predicting Mendelian disease related mutations, its predictions of cancer driver mutations lags behind significantly [29]. Therefore it is unclear if overall biochemical principles behind different disease mutations exist. Our results presented here suggest different types of pathological missense mutations are all governed by the same biochemical principles. In addition, the relevant information required are largely encoded in the protein structure and sequence. Furthermore, they can be effectively extracted by SPRI. We show accurate predictions can be made on cancer driver mutations, even though SPRI was trained using Mendelian disease mutations, requiring no parameter adjustment or re-training using different data.

In addition to structure-derived features, features of evolutionary signals, and biophysical properties are also found to be important for distinguishing deleterious mutations from neutral mutations (see Supplemental Information). Together, they capture essential biological properties whose perturbations contribute to pathogenicity.

A limitation of SPRI is that it is based on analysis of protein structure of monomeric units. Structures of protein complex contains valuable information on protein-protein interactions. As experimental and reliable predicted structures of protein-complexes continue to increase [76, 78, 79] and technical issues [80] being resolved, we anticipate that incorporating knowledge of protein-protein interaction interfaces will further improve models for predictions of pathogenicity of missense mutations.

## 5 Conclusion

Overall, SPRI is highly effective in extracting properties determining pathogenicity encoded in protein structures to provide accurate evaluation of pathological effects of missense mutations and to identify deleterious mutations. SPRI can provide accurate predictions of deleterious missense single mutations of germ line origin associated with Mendelian diseases, and performs favorably when compared to current state-of-the-art methods, as measured by several performance metrics.

The effectiveness of SPRI is transferable and SPRI can be used to identify pathological missense mutations of somatic origin as manifested in complex diseases. This is demonstrated by the success in predicting cancer driver mutations.

In addition, SPRI can discover higher-order spatial units of multiple low recurrence missense mutations, which are strongly indicative of high susceptibility to cancer. Furthermore, SPRI can also predict mutation effects at saturation level, as demonstrated by the example of the copper transport protein.

As SPRI can take advantage of AlphaFold2 predicted structures, it can be deployed for saturation mutation analysis proteome wide and can be used to construct an atlas of effects of saturation mutations. Such an atlas can provide a global view of protein fitness landscape and mutation effects of the whole protein universe.

## 6 Supplement Information

The online version contains supplementary material available at `weblinkdesignatedbythejournal`. Additional files provides information about benchmark datasets, interaction profiles of protein based on alpha shapes, analysis on feature importance, statistics for sequence depth of multiple sequence alignment, ROC and PR curves for each testing fold, performance for different substitutions at the same residue, robustness of structure-derived features using different structures of the same protein, and location preference of deleterious/neutral mutations for well-studied enzymes.

## Availability of data and materials

The pre-computed results on human proteome with known PDB structures can be found at http://sts.bioe.uic.edu/spri/, these benchmark datasets generated and analysed during the current study are available is accessible at https://bitbucket.org/lianglabuic/spri/.

## Authors’ contributions

Conceptualization: BW and JL, with input from YYT. Methodology: BW and JL. Dataset and implementation: BW. Analysis: BW and JL. Additional analysis tools: XL, WT, APR, YYT. Original draft: BW and JL. Review and editing: XL, WT, APR and YYT. Supervision: JL. The authors read and approved the final manuscript.

