## supplemental information for "SPRI: Structure-Based Pathogenicity Relationship Identifier for Predicting Effects of Single Missense Variants and Discovery of Higher-Order Cancer Susceptibility Clusters of Mutations"

### Supplementary Information

#### 1 Technical Details

##### 1.1 Construction of Datasets

Several benchmark datasets have been constructed in previous studies. They are based on the premises that missense mutations known to result in Mendelian disorders and other non-cancer diseases can be regarded as deleterious mutations, whereas mutations with unknown effect can be treated heuristically as neutral mutations. These datasets include HumVar, HumDiv, ExoVar, VariBench, and PredictSNP, and are in wide used [1, 2, 3, 4, 5]. However, these datasets are highly biased, as deleterious mutations and neutral mutations are often from different proteins: there is only a small proportion of proteins possess both deleterious and neutral variants.

Proteins have different degree of deleteriousness. Mutations at sites in functional domains are more likely to have pathological outcomes [6, 7]. In addition, proteins with larger functional surfaces are likely to contain more sites sensitive to substitutions. Hence, proteins with multiple functional domains [8, 9] and larger functional surfaces are less tolerant to mutations, whereas proteins with single function and smaller functional surfaces are more tolerant to mutations.

A significant drawback of using existing datasets is that inconsistent behavior in predictions may result when applied to different types of data, For example, the FATHMM method performs well using the dataset VariBench [10], but exhibits high false positive rate when benchmarking PDB-mapped mutations of the HumDiv dataset [11].

We obtain the HumDiv, HumVar datasets from the Polyphen-2 website `http://genetics.bwh.harvard.edu/pph2/`, and the PredictSNP dataset from `https://loschmidt.chemi.muni.cz/predictsnp/`. We map all mutations to the same version of the canonical protein sequences of Uniprot Jan 2021 release [12]. This reconciles occasional sequence inconsistency among different reference sequences. The human protein sequences and sequence database of eukaryotic reference proteomes are accessed from `https://ftp.uniprot.org`.

We then discard redundant mutations, remove conflicting mutations with both neutral and deleterious labels for the same mutation. Furthermore, we select only those proteins that contain both deleterious and neutral missense mutations. To retrieve the experimental determined Pdb structures, we employ the SeqMapPDB tool to obtain full-coverage structures and partial structural domains, respectively [13, 14]. We then generate the UnifyPDBFull and UnifyPDBAcceptable benchmark datasets. The former contains proteins whose full structures are known, and the latter contains proteins with both full-coverage structures and domains of structures with partial-coverage. The UnifyPDBFull dataset contains 4,231 deleterious variants and 2,791 neutral variants, which are derived from 252 proteins and are mapped to 252 polypeptide chains in PDB structures. The UnifyPDBAcceptable dataset contains 5,999 deleterious variants and 3,485 neutral variants, which are derived from 377 proteins, and are mapped to 444 Pdb structural chains.

We employ the same dataset construction procedure for the proteins having AlphaFold2 predicted structures but no PDB records, and generate the UnifyAF2 dataset. The UnifyAF2 dataset contains 4,980 deleterious and 2,600 neutral variants from 422 proteins. We stratify mutation site residues into three categories of high, medium, and low-confidence level of their predicted structures using the pLDDT score from AlaphFold2. Those of high-confidence have pLDDT above 90, moderate-confidence with pLDDT between 70 and 90, and low-confidence with pLDDT no greater than 70.

#### 1.2 Structure-Derived Features

The structural coordinates of proteins are retrieved from the Pdb database at `https://ftp.rcsb.org/pub/pdb/`. The Alphafold2 predicted structures are obtained from `https://alphafold.ebi.ac.uk`. We calculate the weighted alpha shapes using the van der Waals (VDW) radius of each chemical element, with water molecule probe of 1.4  radius. Atomic interactions are defined by alpha shapes, which captures exact near-neighboring atomic contacts that are dual to Voronoi boundaries separating two inter-residue atoms [15]. We then construct the inter-residue interaction profiles.

For short-range atomic interactions, we define 16 types of element pairs, and vectorize spatial contact information into integer values for each interaction type. We consider only four atom types of C, N, O, and S, distinguishing atoms provided by the mutated residue and its neighboring residue. For example, carbon from the mutation site and nitrogen from the neighboring residue are recorded as *CN*, and carbon from the neighboring residue and nitrogen from the mutation site are recorded as *NC*, so donor and acceptor information is encoded. We dispense with detailed chemical information of orbital hybridisation of chemical element to avoid overfitting [16, 17]. Furthermore, we record only number counts of atomic interactions and ignore their distance or volume overlap measures as proteins often experience conformational fluctuations [18].

We use the Breadth-First Search (BFS) algorithm to obtain atom-interactions at residue level in intermediate- and long-distance range [19]. These are converted to 20-element vectors, where each element represents the number of each amino acid type occurs at the assigned distance range. We use the CASTp server to compute the solvent accessible surface area (SA) and assign property of geometric location for each residue [20]. A salt bridge is considered to exist if the distance between the oxygen atoms in an acidic residue and the nitrogen atoms in a basic residue is within 3.2 [21].

#### 1.3 Encoding Biophysical Changes Introduced by Variants

We regard Asp and Glu as negatively (-1) ionizable residues. Arg, His and Lys as positively (+1) ionizable residues. All remaining amino acids are labelled as neutrally charged (0). For every mutation pattern, we calculate the change of charges by taking the difference of the values of charge labels, which are integer values in the range of -2 to 2.

Changes in atomic composition are also considered, so property changes between the wild-type amino acid and the substituted amino acid residue are incorporated. In most cases, the backbone atomic composition does not change, except when mutation patterns involve Glycine (Gly), which lacks $C_\beta$. This is also duly recorded. For side-chains, we count changes in each of the chemical elements, and obtain an overall atomic compositional change for a given mutation pattern.

#### 1.4 Sequence-Based Features

We employ a standard procedure for multiple sequence alignment (MSA). We first construct the sequence knowledge database from reference proteomes of 1,553 eukaryotic species, excluding that of human. This database contains about

24 millions protein sequences. For each queried human protein, we use BLASTp and Clustal-W2 to obtain its homologs, and select those with identity greater than 30% to the human protein sequence [22, 23]. From the assembled homologous sequences, we construct the MSA using Clustal Omega [24]. We then calculate wild-type frequency, and mutated-type frequency at the mutation site, as well as site-specific entropy, from the aligned MSA. Substitution scores are taken from the BLOSUM62 matrix [25].

#### 1.5 Prediction Model and Performance Comparisons

Overall, the features we use for predictions are structural properties, evolutionary signals, and biophysical properties and changes upon substitutions. All are numerical values, except geometric location, which is categorical. We then train a random forest predictor implemented in R [26]. The number of ensemble trees is set to 500, with each tree fit to a balanced training set such that each tree receives an equal number of positive and negative cases. We use 5-fold cross validation on a stratified test dataset for both UnifyPDBFull and UnifyPDBAcceptable datasets [27]. Prediction results by other methods are retrieved from their open-access web servers or released datasets as of September 18 2021 [28, 10, 2, 29, 1, 30, 31]. Some methods (SPRI, LIST, PMut, Polyphen-2 and Rhapsody) provide a probability value explicitly, others (EVmutation, FATHMM and PROVEAN) provide raw scores, which are scaled to the range between 0 and 1 using min max normalization [32]. We compare different methods by computing Receiver Operating Characteristic (ROC) curves and Precision-Recall (PR) curves. When no predictions can be made by a specific method, we employ an imputation method to assign a random value from uniform distribution in the range of 0 to 1 to compensate the missing predictions [33].

#### 2 Balanced Benchmark Data Sets of UnifyPDBFull and UnifyPDBAcceptable

The HumDiv, HumVar and PredictSNP datasets provide reliable information on deleterious variants contributing towards Mendelian-type disorders. However, only a small number of proteins in each of these datasets contain both neutral variants and deleterious variants as shown in Table 1. Since genes and proteins have different degree of deleteriousness, imbalanced datasets are intrinsically biased.

Table 1: Statistics for Benchmark Datasets in Previous Studies

| | HumDiv | HumVar | PredictSNP |
|---|---|---|---|
| Proteins with Deleterious Variants | 978 | 1,852 | 1,410 |
| Proteins with Neutral Variants | 390 | 8,791 | 10,000 |
| Proteins with Deleterious and Neutral Variants | 390 | 956 | 609 |

To reduce data bias, we construct two new benchmark datasets. We take proteins with variant information in HumDiv, HumVar and PredictSNP, and map them to the same version of canonical protein sequences, with redundant variants removed. We then discard variants with conflicting labels, and include only proteins with both deleterious and neutral variants. We further select proteins with PDB structures by using the SeqMapPDB pipeline [14]. The first resulting dataset UnifyPDBFull contains proteins with full-coverage structures. The second resulting dataset UnifyPDBAcceptable contains proteins with full-coverage and partial-coverage structures, which is a proper superset of UnifyPDBFull. The statistics of the new benchmark datasets are shown in Table 2.

Table 2: Statistics for New Benchmark Datasets Generated in This Study

| | UnifyPDBFull | UnifyPDBAcceptable |
|---|---|---|
| Total number of Proteins | 252 | 377 |
| Number of PDB Monomer Chains | 252 | 444 |
| Proteins with Full-Coverage Structures | 252 | 252 |
| Proteins with Partial-coverage Structural Domains | 0 | 125 |
| Neutral Variants | 2,791 | 3,485 |
| Deleterious Variants | 4,231 | 5,999 |

Note 1: UnifyPDBFull dataset requires the protein to have full-coverage PDB structure, and does not include protein with partial-coverage PDB structure. As a result, the number of proteins with partial-coverage structural domains is 0 on UnifyPDBFull dataset.

Note 2: A protein can have multiple non-overlapping partial structural domains, therefore the number of PDB chains is larger than the number of proteins in the UnifyPDBAcceptable dataset.

Table 3: Dataset Size of Different Methods

| | SPRI UnifyPDBFull | Polyphen-2 HumDIV /HumVar | Rhapsody 2020 Integrated | FATHMM | PMut | LIST |
|---|---|---|---|---|---|---|
| Size of Dataset | 7,022 | 9,476 /21,978 | 27,655 | 49,414 | 65,281 | 10,208 |

### 3 Interaction Profiles of Protein Based on Alpha Shapes

We use the structure of chain `A` of protein PDB `6RTX` as an example to illustrate the construction of interaction profiles. The computation of its interaction profile using alpha shapes is shown in Fig 1. Here a vertex represents the center of a heavy atom, and an edge represents the connection between two atoms whose Voronoi regions intersect with each other and the intersection is within the union of atoms. The interaction profiles extracted from vertexes-connecting edges can provide us with detailed pairs of atomic element in short-range, intermediate-, and long-range interaction, according to their relative distances to the mutation site residue.

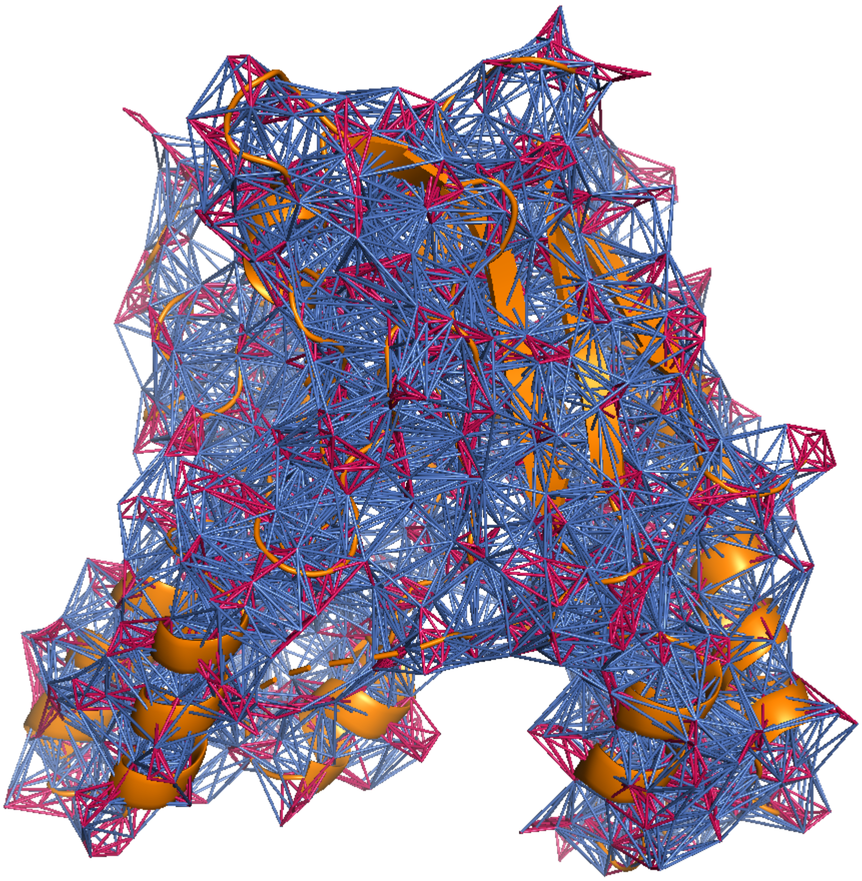

Figure 1: The alpha edges used for computing the interaction profiles of protein structure for `6RTX` chain `A`. The orange ribbons represent the carton model of the protein. Red edges are intra-residue atomic interaction within the same residue. Blue edges are inter-residue atomic interaction between different residues.

### 4 Analysis of Feature Importance

To identify which features contribute most towards accurate prediction, we report the variable importance plot, in which features with high-ranked mean decrease in Gini coefficient are considered to be more important for predictions [34].

The importance plot shows that features from all of three categories (structural properties, evolutionary signals from sequence analysis, and biophysical properties and changes upon substitutions) play important roles in distinguishing deleterious mutations from neutral mutations. Important features extracted from protein structures include solvent accessible surface area (SA), short-range atomic interactions such as carbon-carbon interaction recorded as *CC*, nitrogen-carbon interaction (*NC*), and oxygen-carbon interaction (*OC*), the residue interaction profiles in intermediate-range (*e.g.* Leu_L2, Val_L2) and long-range (*e.g.* Leu_L3, Phe_L3). The biophysical changes upon substitutions also provide important features. These includes the change in charge, changes in side-chain atoms (*e.g.* carbon, nitrogen). Furthermore, all features extracted from protein-specific alignments are highly ranked, including frequency of mutated type of residue, and frequency of wild-type residue at the mutation site, BLOSUM score, and the site-specific evolutionary entropy.

We also use different subsets of features to understand effectiveness of evolutionary, structural and biophysical features. From the ROC curves and Precision-Recall curves, we can conclude that the model trained on structural+biophysical (non-evolutionary) features exhibits similar performance as the model using evolutionary features only, while none of the subsets of features is capable to achieve overall performance as the model trained on full features.

The results from this analysis suggest that features extracted from the three different categories all represent essential biological characteristics underlying the effects of substitutions, and they complement each other to provide fine-grained information for each specific mutation pattern, enabling reliable prediction.

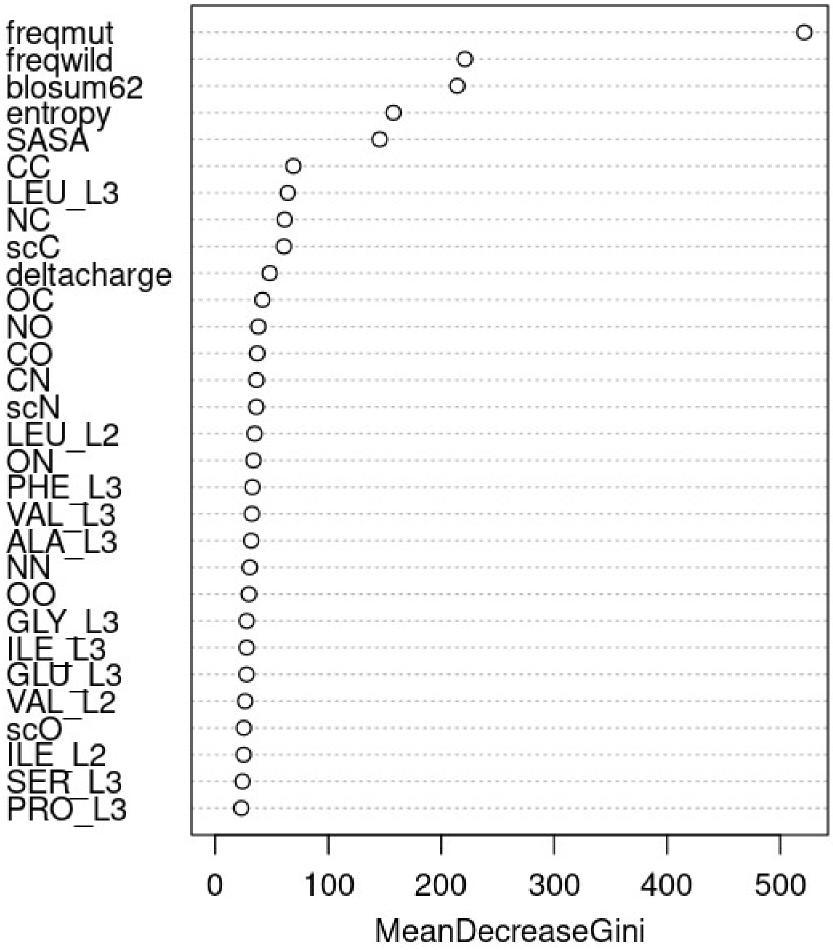


Figure 2: Plot of high-ranked variable importance. The higher value of mean decrease gini coefficient indicates the corresponding feature is more important to contribute the prediction performance of the model.

Table 4: Average performance of sub feature sets on UnifyPDBFull dataset

| | Full | Structural+Biophysical | Evolutionary | Structural | Biophysical |
|---|---|---|---|---|---|
| Average AUROC | 0.940 | 0.879 | 0.891 | 0.845 | 0.754 |
| Average AUPRC | 0.956 | 0.917 | 0.913 | 0.891 | 0.575 |

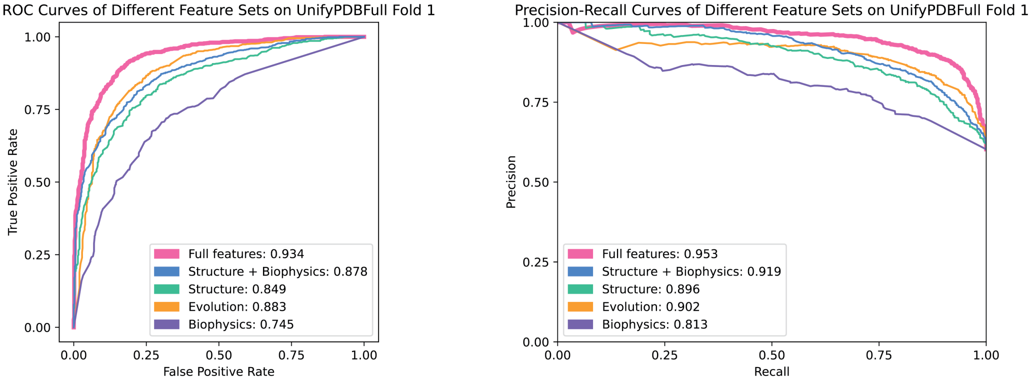


Figure 3: Performance of Different Subsets of Features on UnifyPDBFull Fold 1

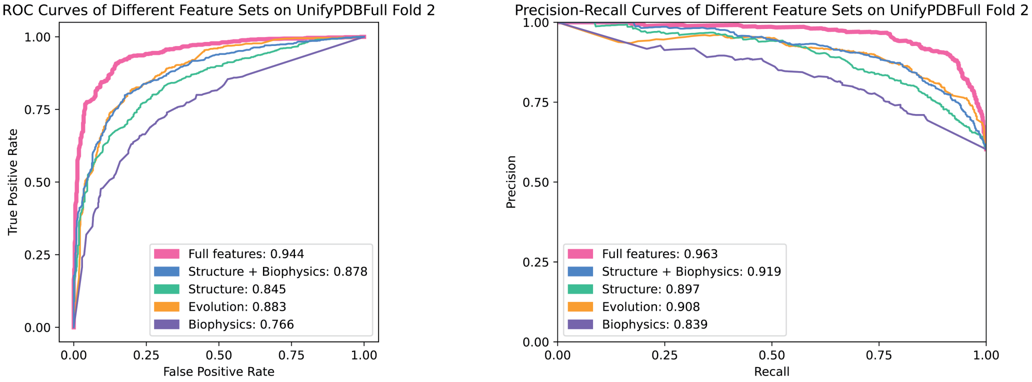


Figure 4: Performance of Different Subsets of Features on UnifyPDBFull Fold 2

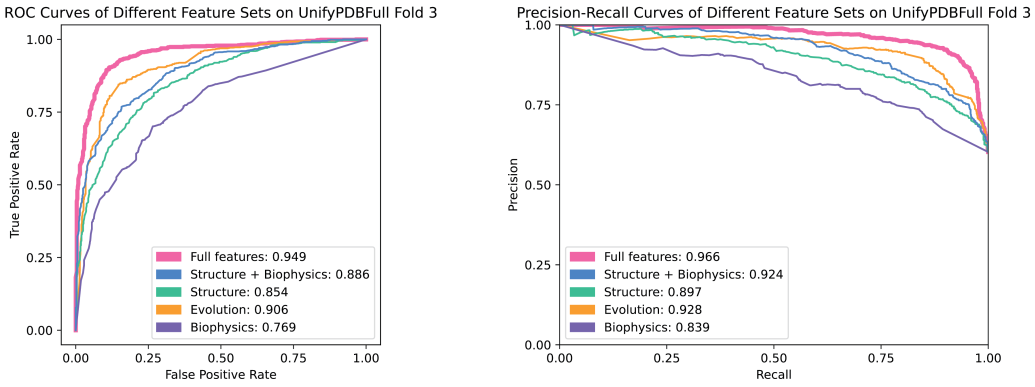


Figure 5: Performance of Different Subsets of Features on UnifyPDBFull Fold 3

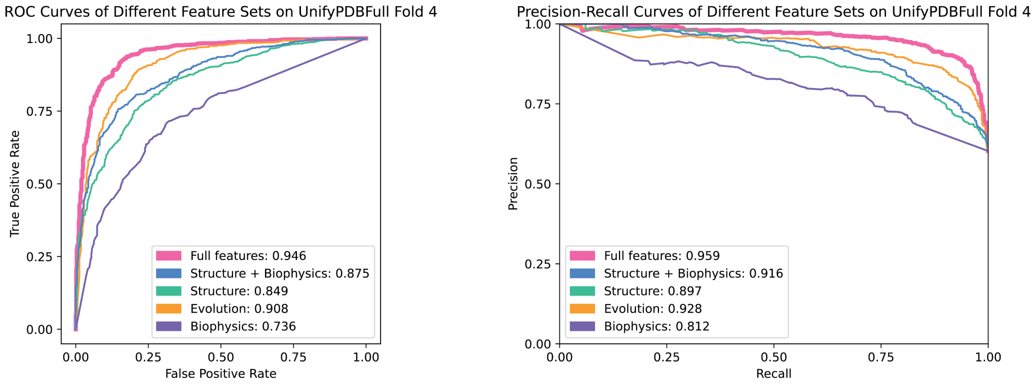


Figure 6: Performance of Different Subsets of Features on UnifyPDBFull Fold 4

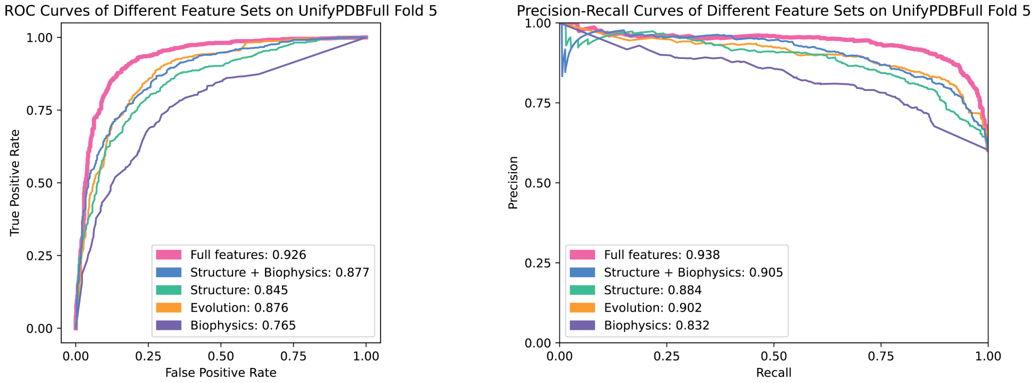


Figure 7: Performance of Different Subsets of Features on UnifyPDBFull Fold 5

### 5 Depth of Multiple Sequence Alignment (MSA)

The depth of multiple sequence alignment is the total number of individual sequences aligned in a protein-specific MSA. Overall, most of 377 protein-specific MSAs in this study have adequate alignment depth that is above 50. Specifically, 94.4% (356 out of 377) MSAs have an alignment depth of at least 100 individual sequences. The median of the alignment depth is 404, the first quartile is 226, and the third quartile is 496. These alignment depths indicate that the evolutionary signals can be reliably estimated from MSA.

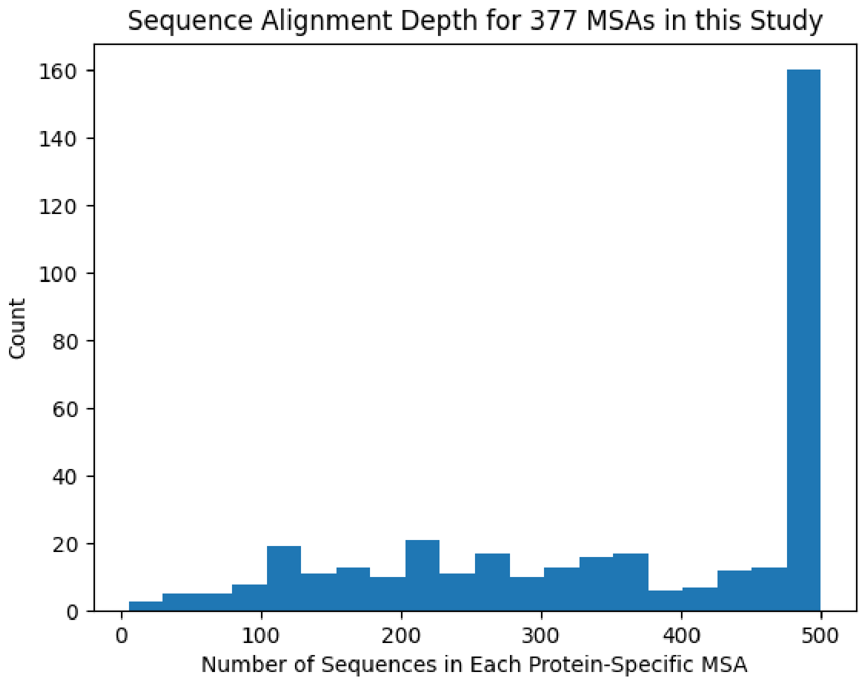


Figure 8: Distribution of sequence alignment depth

### 6 Performance on UnifyPDBFull dataset by Each Fold with Imputation of Missing Cases

We report the Receiver operating characteristic (ROC) and Precision-Recall (PR) curves for each fold of the UnifyPDBFull stratified cross validation. We use default thresholds provided by FATHMM, PMut, Polyphen-2, PROVEAN and Rhapsody to distinguish deleterious mutations from neutral mutations [10, 29, 1, 30]. We use the optimal thresholds for EVmutation, LIST, and our method at the highest Matthews correlation coefficient (MCC) values for evaluation.

Most of the 8 methods we tested have good coverage (>0.980) on UnifyPDBFull dataset, except EVmutation and Rhapsody, which have a coverage of 0.740 and 0.930, respectively. To account for missing predictions so a fair comparison can be made, we employ an imputation process which assigns a random value generated from the uniform distribution from 0 to 1 for any missing prediction.

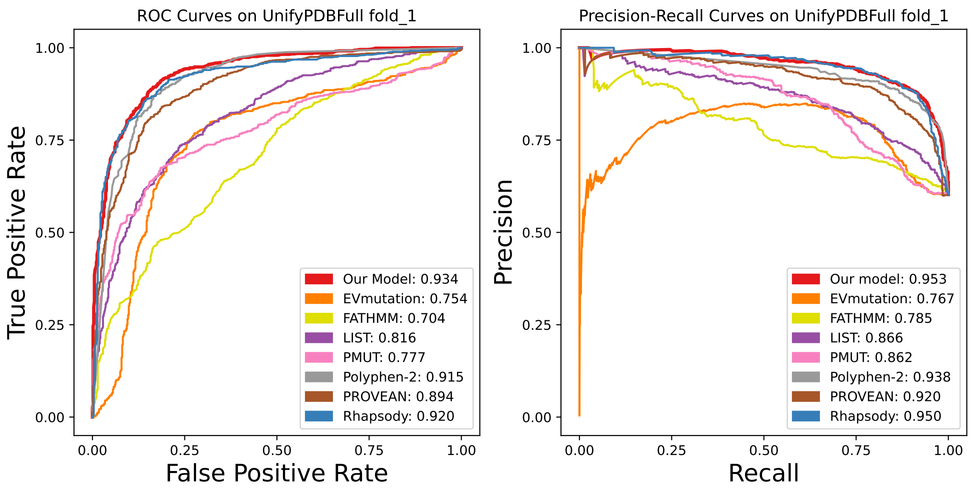


Figure 9: Performance on UnifyPDBFull Fold 1 with imputation of missing predictions

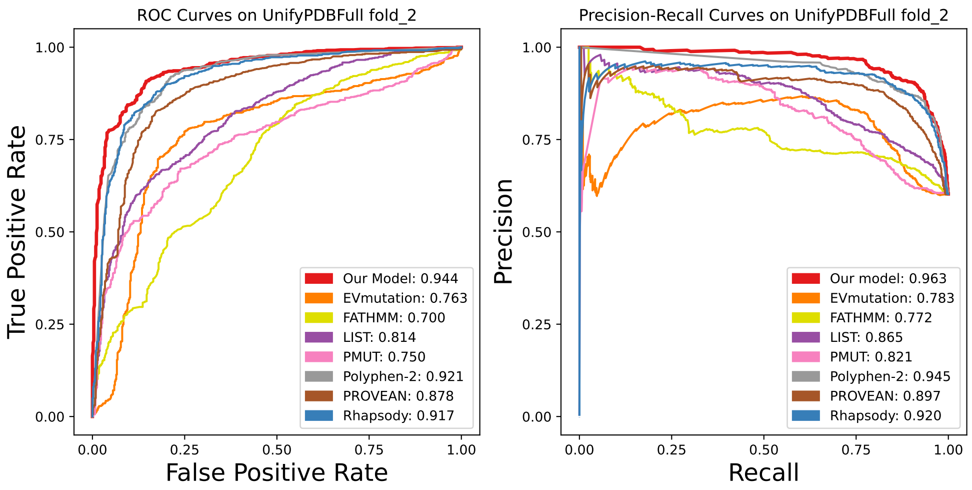


Figure 10: Performance on UnifyPDBFull Fold 2 with imputation of missing predictions

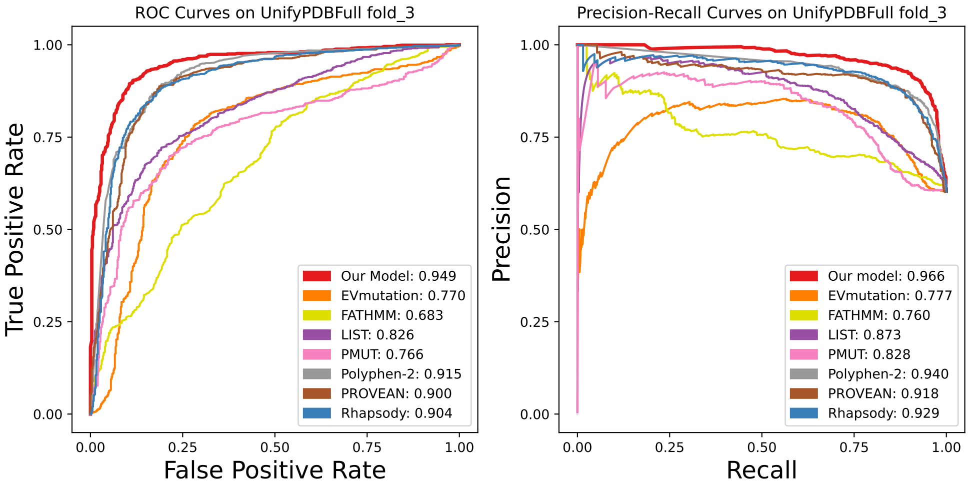


Figure 11: Performance on UnifyPDBFull Fold 3 with imputation of missing predictions

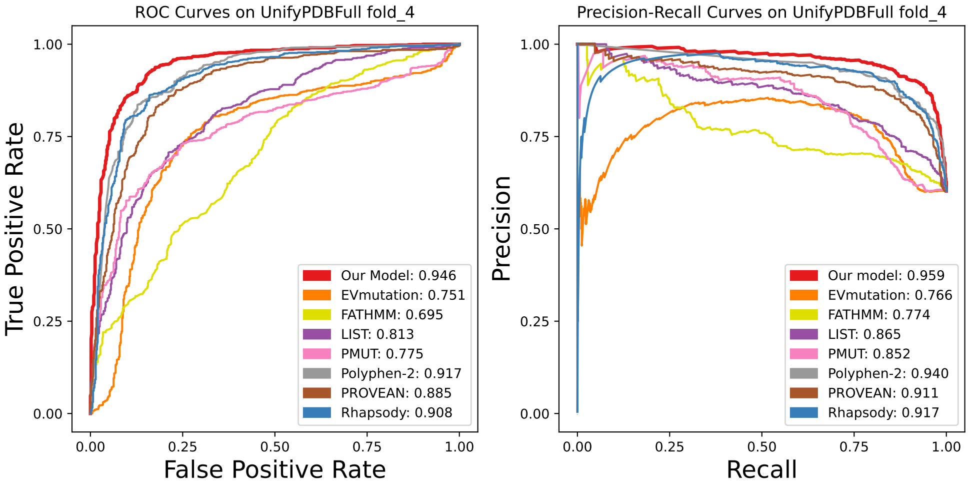


Figure 12: Performance on UnifyPDBFull Fold 4 with imputation of missing predictions

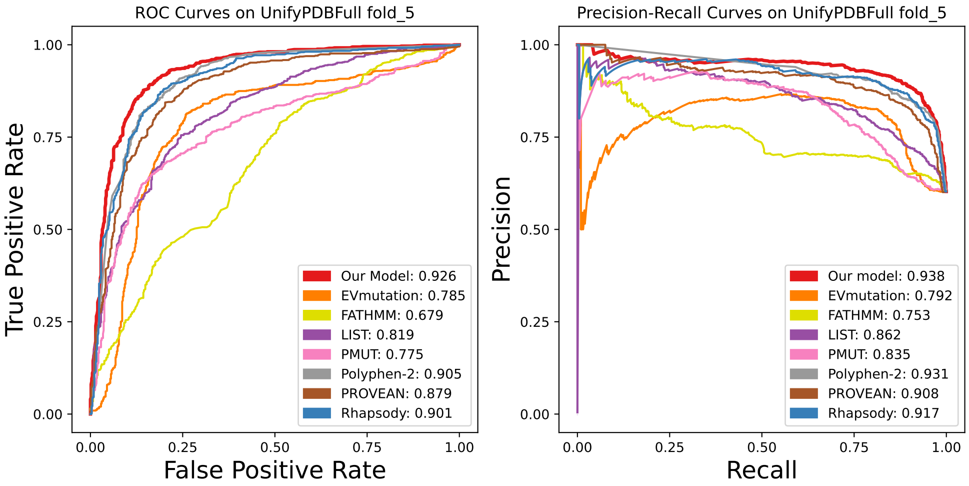


Figure 13: Performance on UnifyPDBFull Fold 5 with imputation of missing predictions

Table 5: Completeness/Coverage for Each Method in Each Fold

| SPRI | EVmutation | FATHMM | LIST | PMUT | Polyphen-2 | PROVEAN | Rhapsody |
|---|---|---|---|---|---|---|---|
| 1.0 | 0.744 | 0.986 | 0.986 | 0.992 | 0.988 | 0.986 | 0.927 |
| 1.0 | 0.731 | 0.982 | 0.982 | 0.989 | 0.982 | 0.982 | 0.943 |
| 1.0 | 0.745 | 0.990 | 0.993 | 0.986 | 0.992 | 0.990 | 0.922 |
| 1.0 | 0.737 | 0.989 | 0.988 | 0.992 | 0.985 | 0.989 | 0.929 |
| 1.0 | 0.741 | 0.989 | 0.986 | 0.989 | 0.990 | 0.989 | 0.929 |

### 7 Performance on UnifyPDBAcceptable dataset by Each Fold with Imputation of Missing Cases

We report the ROC and Precision-Recall curves for each fold of UnifyPDBAcceptable stratified cross validation. Both ROC curves and PR curves are used to examine performance for binary classification models, with PR curves providing a more informative assessment of imbalanced dataset.

Most of the 8 methods have good coverage ($> 0.980$) on UnifyPDBAcceptable dataset, except EVmutation and Rhapsody which have coverage of 0.711 and 0.941, respectively. To account for missing predictions so a fair comparison can be made, we employ an imputation process which assigns a random value generated from the uniform distribution from 0 to 1 for any missing prediction.

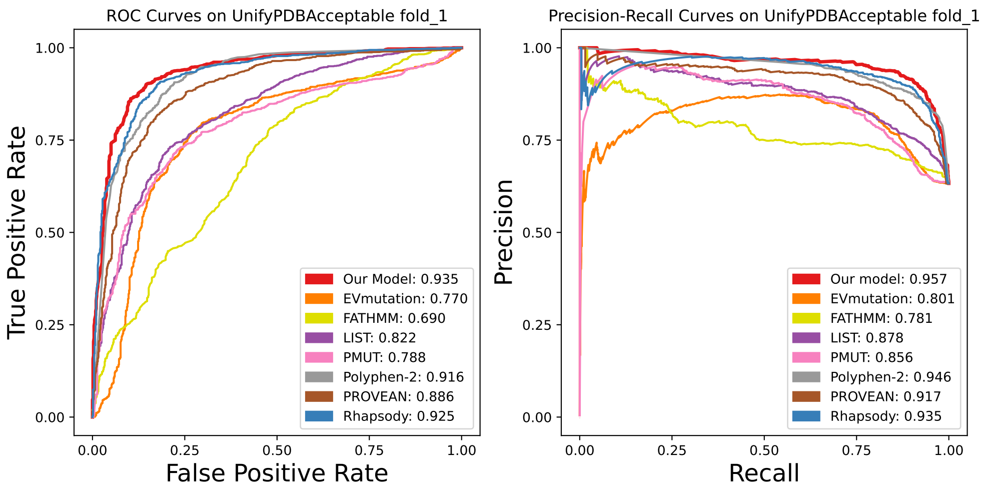


Figure 14: Performance on UnifyPDBAcceptable Fold 1 with imputation of missing predictions

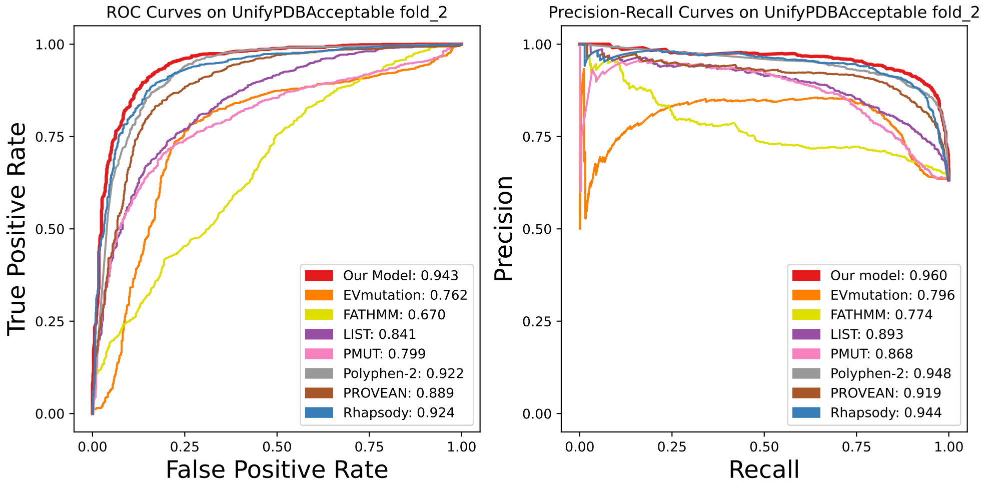


Figure 15: Performance on UnifyPDBAcceptable Fold 2 with imputation of missing predictions

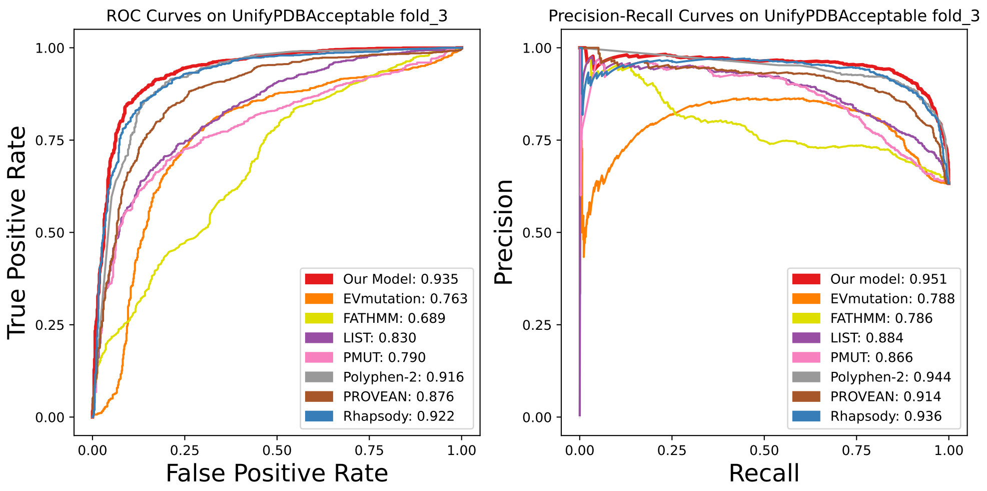


Figure 16: Performance on UnifyPDBAcceptable Fold 3 with imputation of missing predictions

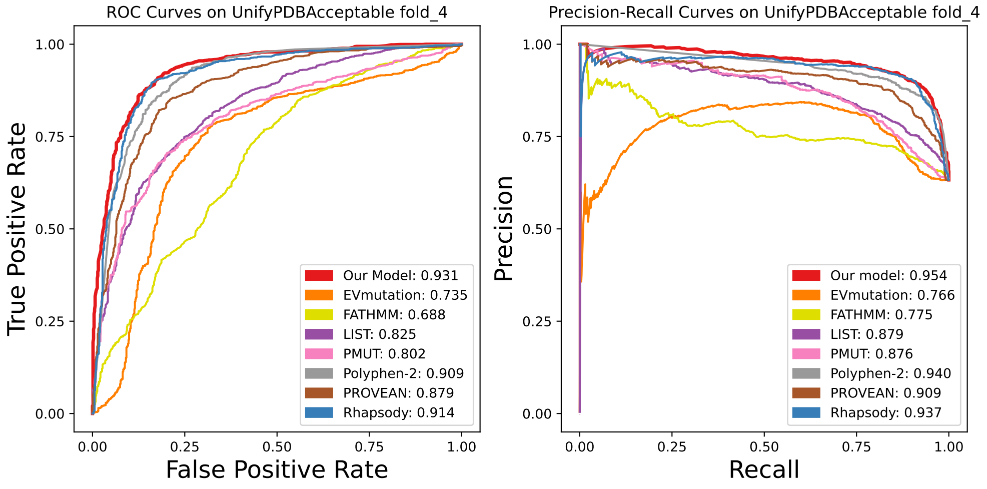


Figure 17: Performance on UnifyPDBAcceptable Fold 4 with imputation of missing predictions

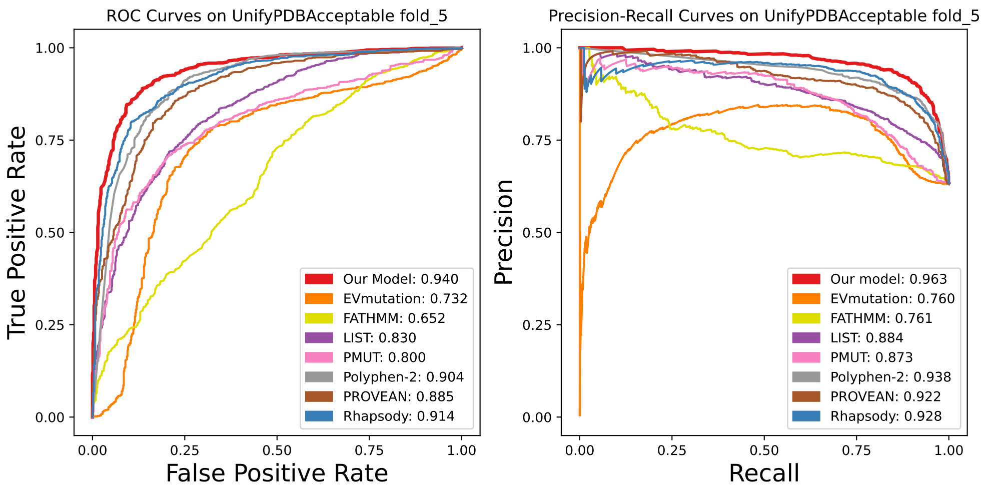


Figure 18: Performance on UnifyPDBAcceptable Fold 5 with imputation of missing predictions

Table 6: Completeness/Coverage for Each Method in Each Fold

| SPRI | EVmutation | FATHMM | LIST | PMUT | Polyphen-2 | PROVEAN | Rhapsody |
|---|---|---|---|---|---|---|---|
| 1.0 | 0.699 | 0.991 | 0.989 | 0.988 | 0.989 | 0.991 | 0.941 |
| 1.0 | 0.707 | 0.992 | 0.992 | 0.986 | 0.989 | 0.992 | 0.948 |
| 1.0 | 0.719 | 0.991 | 0.991 | 0.992 | 0.991 | 0.991 | 0.939 |
| 1.0 | 0.709 | 0.994 | 0.993 | 0.987 | 0.992 | 0.994 | 0.944 |
| 1.0 | 0.722 | 0.985 | 0.986 | 0.982 | 0.987 | 0.985 | 0.934 |

Table 7: Prediction for Variants Occurring at the Same Mutation Sites of Human Ornithine Carbamoyltransferase

| **Truth** | **Index** | **Wild-type** | **Mutant** | **Deleterious Probability** | **Prediction** | **Correctness** |
|---|---|---|---|---|---|---|
| Deleterious | 56 | M | T | 0.796 | Deleterious | Correct |
| Neutral | 56 | M | I | 0.208 | Neutral | Correct |
| Deleterious | 125 | T | M | 0.160 | Neutral | Incorrect |
| Neutral | 125 | T | K | 0.278 | Neutral | Correct |
| Deleterious | 159 | I | T | 0.796 | Deleterious | Correct |
| Neutral | 159 | I | V | 0.186 | Neutral | Correct |
| Deleterious | 160 | I | S | 0.798 | Deleterious | Correct |
| Neutral | 160 | I | V | 0.126 | Neutral | Correct |
| Deleterious | 255 | H | P | 0.554 | Deleterious | Correct |
| Neutral | 255 | H | C | 0.168 | Neutral | Correct |
| Neutral | 255 | H | R | 0.028 | Neutral | Correct |
| Deleterious | 320 | R | L | 0.890 | Deleterious | Correct |
| Neutral | 320 | R | Q | 0.476 | Neutral | Marginally Correct |

### 8 Evaluating Mutation Effects of Different Substitutions at Same Site

Each residue site in a protein can in principle have 19 different substitutions, and they may introduce different pathological effects. We have tested how well SPRI can distinguish different mutation effects from different substitutions occurring at the same mutation site.

In the UnifyPDBFull dataset, human ornithine carbamoyltransferase (OTCase, uniprot: P00480) has the most number of known substitutions at the same sites. 6 mutation site residues have both deleterious and neutral mutations. To ensure accurate assessment of prediction results, we carried out a Leave-One-Protein-Out validation test, in which variant of OTCase is not included in the training process. Our model predicts the effects of 12 out of 13 variants correctly. The results suggest that our model is capable to provide accurate pathological effects for different substitutions at the same mutation site.

### 9 Robustness in Predictions Using Different Structures

To verify that our method can provide robust predictions for mutations when different structures are used, we examine predictions made using different PDB structures of the same protein, each capturing the protein in a particular conformation from a fluctuating ensemble. We select the protein uroporphyrinogen decarboxylase for this task. The uroporphyrinogen decarboxylase (UniProt: P06132) has five X-ray resolved conformations in the PDB, namely, 1R3Q chain A, 1R3Y chain A, 3GVR chain A, 1URO chain A, and 3GVQ chain A.

We choose chain A of 1URO as the reference structure, and compare its property values and its predicted pathological effects with those using other PDB structures. In the UnifyPDBFull dataset, this protein has 32 deleterious mutations on 30 different mutation sites (L20P, G25E, W34R, D79N, A80S, F84I, V134Q, T141I, R144P, P150L, P150Q, P150S, G156D, T160I, M165R, E167K, G168R, Y182C, S188R, L192P, R193P, G205R, E218K, F232L, L253Q, V256G, I260T, W284R, K297N, D306Y, N336S, H349D), and 8 neutral mutations on 8 mutation sites (L68P, P106L, R113T, R120C, G132D, V208A, N230S, G303V). Our task is to compare structural properties for 38 mutation sites on 4 alternative structures with the reference structure.

Specifically, we compare for each mutation site a total of 58 numerical features (16 types of atomic interactions, 20 types of intermediate-range residue interactions, 20 types of long-range residue interactions, surface area, and salt bridge) and 1 categorical feature (geometric location). We then compare 8,968 pairs of features ( 59 features $\times$38 mutation sites $\times$ 4 pairs of alternative structures and reference structure). Among 81.24% (7,286 out of the 8,968) cases, the integer and categorical values of structure-derived features are exactly the same as their paired values computed based on the reference structures. An additional 4.66% (418 out of 8,968) value pairs are within 20% integer or real values of those obtained using reference structure.

We have also compared the predicted pathological effects of mutations using different structures. In total, among the 160 prediction pairs (40 mutations $\times$ 4 pairs of alternative structures and reference structure), the predictions all give the same classification label, *i.e.* deleterious versus neutral. Furthermore, the difference of probability values for a specific mutation between the reference structure and an alternative structures has a mean value of 0.018 and standard deviation of 0.016.

Overall, these results demonstrate that our method is robust in computing structural properties for a protein when using different structures in different bound states, or resulting from different experimental conditions.

#### 10 Statistics for Location Preference of Deleterious/Neutral Mutations

We provide detailed statistics of analysis of the spatial relationship of deleterious and neutral mutations in the catalytic and neighboring regions for 19 well-studies enzymes in the UnifyPDBFull dataset. Here the null hypothesis is that deleterious and neutral mutations exhibit the same likelihood to be located in catalytic regions or neighboring regions, and the alternative hypothesis is that deleterious variants are more preferably located at the catalytic region and its neighboring region, while neutral mutations are more likely to occur at the outside region. The Fisher's exact test results in a $p$-value of $4.2\times10^{-14}$, which strongly rejects the null hypothesis.

Table 8: Location Preference

| Location | Deleterious Variants | Neutral Variants |
|---|---|---|
| Catalytic and Neighboring Region | 305 | 34 |
| Outside Region | 224 | 113 |
| $P$-value | $4.2\times10^{-14}$ | |

### 11 Threshold Settings for patHOS

Among the 200,402 HOSs on 2,712 proteins in the cancer mutation data, only 0.387% HOSs have $\geq 5$ deleterious mutations, and only 4.358% HOSs have the normalized pathogenicity score $\theta \geq 0.05$ (Fig SI 19-20). As both metrics are informative and complement each other, we therefore take the thresholds of $\geq 5$ deleterious mutations and $\theta \geq 0.05$ to control false positives. This approach works well, as only 0.283% HOSs exceed this dual thresholds.

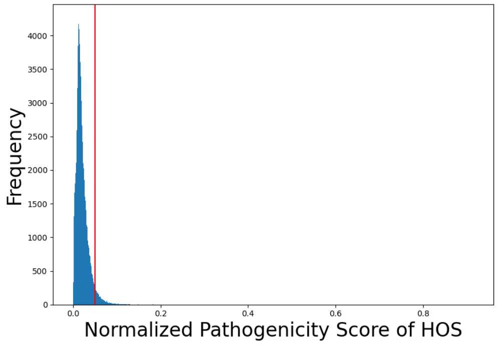


Figure 19: Histogram Plot of Normalized Pathogenicity Score

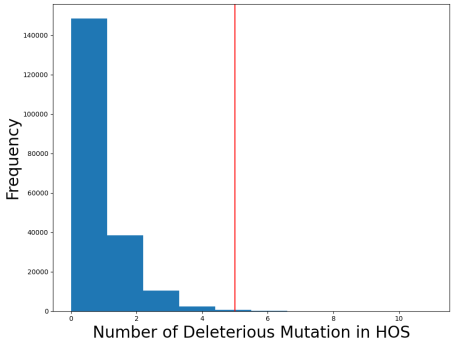


Figure 20: Histogram Plot of Number of Deleterious Mutations in HOSs

caption

#### 12 Literature Evidence for Novel Cancer Genes Identified by SPRI

SPRI identifies 29 novel genes that exhibits two individual patHOSs, which have strong susceptibility in tumorigenesis. Among these, 22 out of 29 genes have literature evidence which supports their relevance with cancer. Details are listed below.

**List 1. literature and cancer type information for these novel driver genes identified by** SPRI

| Genes | Literatures | Cancer Types |
|---|---|---|
| BCHE | PMID: 35915658<br>PMID: 32998690<br>PMID: 34486544 | Endometrial cancer<br>Lung adenocarcinoma<br>Neuroblastoma |
| SLC14A1 | PMID: 36257969<br>PMID: 37004669<br>PMID: 21750109 | Prostate cancer<br>Renal cancer<br>Urinary bladder cancer |
| AOX1 | PMID: 31383940<br>PMID: 30289108 | Bladder cancer<br>Prostate cancer |
| CHAT | Not available | Not available |
| PCDH10 | PMID: 33271263<br>PMID: 22206871 | Colorectal cancer<br>Gastric cancer |
| DPYS | PMID: 25193387<br>PMID: 18446232 | Prostate cancer<br>Colon cancer<br>Breast cancer |

| | | |
|---|---|---|
| MME | Not available | Not available |
| GLUD2 | PMID: 30314897<br>PMID: 25225364 | Glioblastoma |
| ENPP2 | PMID: 35409077<br>PMID: 34769391 | Breast cancer |
| PLG | PMID: 36376702<br>PMID: 30542696<br>PMID: 34745945 | Gastric cancer |
| ACSM2A | Not available | Not available |
| KCNA2 | Not available | Not available |
| ALPP | PMID: 33356330<br>PMID: 35406529 | Ovarian cancer |
| C6 | Not available | Not available |
| F2 | PMID: 30542696 | Colorectal cancer |
| ACTB | PMID: 32795414<br>PMID: 34486492<br>PMID: 23266771 | pan-cancer |
| GABRA1 | PMID: 22038115<br>PMID: 34001135 | Colorectal cancer<br>Glioma |
| POSTN | PMID: 33317907<br>PMID: 34140509<br>PMID: 31968254 | Ovarian cancer<br>Human fibrotic skin diseases<br>Colorectal cancer |
| XDH | PMID: 29551504<br>PMID: 34496841 | Gastric cancer<br>Liver cancer<br>Breast cancer |
| PRMT8 | Not available | Not available |

| | | |
|---|---|---|
| ATP11C | PMID: 24371231<br>PMID: 24904167 | Lung cancer |
| RALA | PMID: 35626682<br>PMID: 34118960<br>PMID: 29047224 | Pan-cancer<br>Breast cancer |
| GNAO1 | PMID: 34900204<br>PMID: 24366063<br>PMID: 32898863<br>PMID: 21317923 | Hepatocellular carcinoma<br>Colorectal cancer<br>Lymphoblastic leukemia<br>Breast cancer |
| AMY2B | Not available | Not available |
| EEF1A2 | PMID: 33473168<br>PMID: 28923030<br>PMID: 14588074 | Lung cancer<br>Prostate cancer<br>Ovarian cancer |
| PNLIP | PMID: 23918603<br>PMID: 36631248 | Pancreatic cancer<br>Pancreatitis (non cancer) |
| DPYSL3 | PMID: 30498031<br>PMID: 29514686<br>PMID: 34691191 | Breast cancer<br>Lung cancer<br>Colon cancer |
| CYP11B1 | PMID: 29542002<br>PMID: 35139664<br>PMID: 34821221 | Adrenal cortical carcinoma<br>Aldosteronism (majority are not tumor) |
| ST8SIA3 | PMID: 31466399<br>PMID: 26794090<br>PMID: 35832124 | Glioblastoma<br>Endometrial cancer |

### 13 Performance Comparison on Likely Cancer Passenger Mutations

The current practice in many studies treats mutations with allele frequency above a threshold (typically, MAF$>0.01$) in the general population or a random mutation from statistical sampling on nucleotide changes to be neutral or passenger mutations [35, 36, 37]. This is somewhat problematic, as passenger mutations should be actual observed mutations existing in cancer patients. A recent study integrating enrichment in linear sequence, in 3D structures, and in function annotation of a specific mutation demonstrated strong predictability of driver mutations [38]. We therefore take the approach that if a mutation lacks significant enrichment signals in both linear sequence and in 3D environment, it is likely a passenger mutation.

#### 13.1 Data set of likely cancer passenger mutations

As currently there is no clearly defined neutral data set of passenger mutations in tumorigenesis that is widely agreed upon, we develop a new data set of neutral passenger mutations based on the following criteria: 1) the mutation does not appear only in the general population, but must be observed in collected cancer samples as recorded in the COSMIC database, 2) the mutation can be mapped to a high quality experimentally determined PDB structure, 3) the mutation must not occur on a known cancer driver gene as defined by the CGC list, 4) the gene should harbor $\leq 10$ Mutations, which suggests such a gene is likely a background gene in cancer, 5) the specified mutation should have no more than 2 recurrences in COSMIC pan-cancer data, 5) the HOS anchored at this mutation should have a few observed ($<3$) co-clustering mutations collected from cancer, to ensure that it is likely an incidental mutation located at this HOS. We take the same cancer mutation dataset in constructing patHOS (239,164 mutations from 200,402 unique mutation site residues on 2,712 proteins), and obtain 117 mutations as passenger mutations in 31 proteins. We recognize this neutral set is not ground truth, but is the best we can obtain as exactly constructed experimental and clinical data are currently not yet available.

Table 10: Performance on Likely Cancer Passenger Mutations

| | SPRI | EVMUTATION | FATHMM | LIST | POLYPHEN-2 |
|---|---|---|---|---|---|
| True Negative (predicted as neutral) | **73** | 12 | 62 | 63 | 55 |
| False Positive (predicted as deleterious) | 44 | 9 | 8 | 49 | 61 |
| No prediction | 0 | 96 | 47 | 5 | 1 |

### 14 Performance on UnifyAF2 Dataset

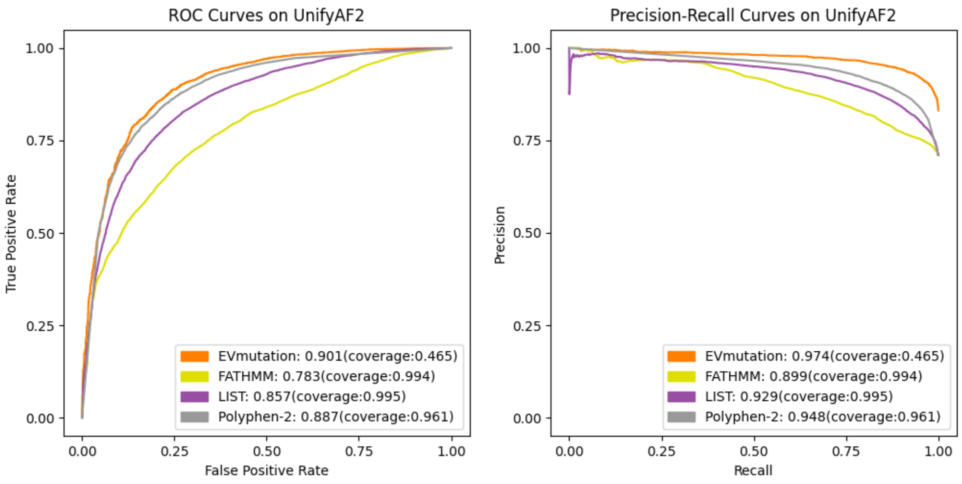


Figure 21: Performance on UnifyAF2 dataset without imputation of missing predictions (coverage revealed)

Table 11: Performance on UnifyAF2 dataset

| | SPRI on High-confidence | SPRI on Moderate-confidence | SPRI on Low-confidence | EVmutation | FATHMM | LIST | Polyphen-2 |
|---|---|---|---|---|---|---|---|
| Completeness (coverage) | 1 | 1 | 1 | 0.465 | 0.994 | 0.995 | 0.961 |
| Recall | **0.895** | 0.866 | 0.608 | 0.881 | 0.903 | 0.661 | 0.841 |
| Specificity | 0.770 | 0.798 | **0.889** | 0.759 | 0.573 | 0.876 | 0.782 |
| Precision | 0.916 | 0.917 | 0.796 | **0.948** | 0.822 | 0.929 | 0.905 |
| Accuracy | **0.862** | 0.847 | 0.773 | 0.860 | 0.736 | 0.723 | 0.824 |
| F-1 score | 0.905 | 0.890 | 0.689 | **0.913** | 0.812 | 0.772 | 0.872 |
| MCC Score | **0.653** | 0.639 | 0.527 | 0.572 | 0.370 | 0.487 | 0.596 |
| AUROC | **0.908** | 0.905 | 0.854 | 0.901 | 0.783 | 0.857 | 0.887 |
| AUPRC | 0.956 | 0.950 | 0.810 | **0.974** | 0.889 | 0.929 | 0.948 |

#### 15 Specified Data Used in Model Training, Testing and Validation

In the case studies of glutathione synthetase (one entry of 19 M-CSA enzymes), we used the subset of UnifyPDBFull, where all the 19 M-CSA enzymes were left out. The model was trained with the remaining 233 proteins, and tested on the glutathione synthetase and other 18 enzymes.

For the 19 enzymes from MCSA, we used the subset of UnifyPDBFull, where all the 19 M-CSA enzymes were left out. The model was trained with the remaining 233 proteins, and tested on the 19 M-CSA enzymes.

For the proteins in the patHOS study, we use the model trained on the whole UnifyPDBFull dataset, and employ it to predict the large-scale 239,164 cancer mutations in 2,712 proteins. As only 611 out of 239,164 (0.25%) cancer mutations appear in the UnifyPDBFull training dataset, 99.75% of cancer mutations are not seen by the SPRI predictor.

As ATOX1 (uniprot: O00244) is not in UnifyPDBFull dataset, our model was trained on the whole UnifyPDBFull dataset to predict mutation saturation of ATOX1.

For the cancer mutation predictions of 26 proteins, we used a subset of 247 proteins from the UnifyPDBFull set after excluding the 5 CGC genes. The performance is then tested on the 26 CGC genes, none of which were in the training set.

For the UnifyAF2 study, UnifyAF2 contains proteins without high-quality PDB entries. None of the mutations in UnifyAF2 is included in either UnifyPDBFull or UnifyPDBAccpetable data sets. Our model was trained on the whole UnifyPDBFull dataset, and tested on UnifyAF2.

The ROC and Precision-Recall curves using the UnifyPDBFull dataset (subplots of a and b in main text Figure 3) are benchmarked on one fold test dataset. The performance shown in main text Table 1 are the average values of 5 test datasets of a 5-fold cross validation test. More details on ROC and Precision-Recall curves of each fold of the 5-fold cross validation is in SI Section 6.

The ROC and Precision-Recall curves using UnifyPDBAccptable dataset (subplots of c and d in main text Figure 3) are benchmarked on one fold test dataset. The performance shown in main text Table 2 are the average values of 5 test datasets datasets of a 5-fold cross validation test. More details on ROC and Precision-Recall curves of each fold of the 5-fold cross validation is in SI Section 7.
